# A robust receptive field code for optic flow detection and decomposition during self-motion

**DOI:** 10.1101/2021.10.06.463330

**Authors:** Yue Zhang, Ruoyu Huang, Wiebke Nörenberg, Aristides B. Arrenberg

**Author notes:** **Contact info**, Twitter: @ArrenbergLab.

## Abstract

The perception of optic flow is essential for any visually guided behaviour of a moving animal. To mechanistically predict behaviour and understand the emergence of self-motion perception in vertebrate brains, it is essential to systematically characterize the motion receptive fields (RFs) of optic flow processing neurons. Here, we present the fine-scale RFs of thousands of motion-sensitive neurons studied in the diencephalon and the midbrain of zebrafish. We found neurons that serve as linear filters and robustly encode directional and speed information of translation-induced optic flow. These neurons are topographically arranged in pretectum according to translation direction. The unambiguous encoding of translation enables the decomposition of translational and rotational self-motion information from mixed optic flow. In behavioural experiments, we successfully demonstrated the predicted decomposition in the optokinetic and optomotor responses. Together, our study reveals the algorithm and the neural implementation for self-motion estimation in a vertebrate visual system.

## Introduction

When you walk across a street, the image of the street is also shifted across your retina. This global retinal image shift induced by translational and/or rotational self-motion is known as optic flow ^1–3^. Optic flow is used for self-motion estimation ^4–8^, which is a critical task for all visually guided navigation, as well as for gaze and posture stabilization behaviours (Figure 1A).

**Figure 1.**
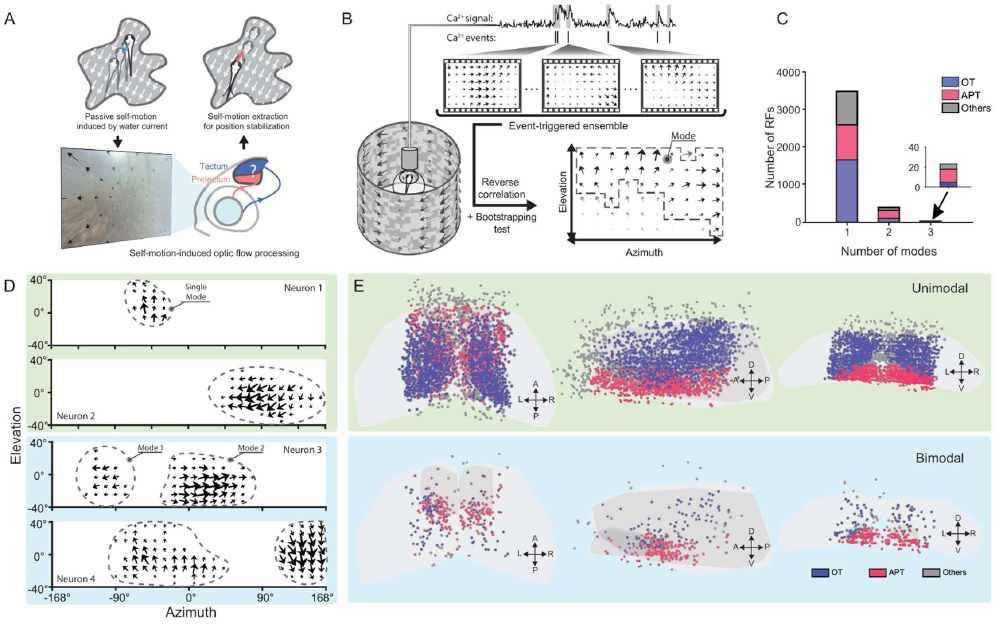
Characterization of unimodal receptive fields (RFs). (A) Illustration of the challenge of extracting self-motion from optic flow for visually guided self-stabilization behaviour (B) Illustration of the RF estimation method using a contiguous motion noise stimulus (see also STAR Methods and Figure S1). (C) Histogram of RF mode number for neurons in the optic tectum (OT), area pretectalis (APT) and adjacent brain regions. (D) Examples of two unimodal (green) and two bimodal motion RFs (blue). The direction and size of the arrows indicate the local preferred direction and the motion sensitivity within the RF, see also Figure S1. (E) Anatomical location of neurons with unimodal RFs (n=3504, in 7 fish) and bimodal RFs (n = 398). The anterior-posterior (AP), dorsal-ventral (DV) and left-right (LR) axes are labelled.

The anatomical substrates associated with this task have been identified in a range of species. For instance, optic flow is processed in the medial superior temporal and parietal cortical areas in primates ^8–13^, the nucleus lentiformis mesencephali in pigeons (Wylie, 2000), and the lobula plate in flies ^17^. In zebrafish, the pretectal area processes optic flow and is necessary for body position and gaze stabilization in optomotor responses (OMR) and optokinetic responses (OKR), respectively ^18–21^.

Knowing which neurons are involved does not answer how these neurons process optic flow and mediate stabilization behaviour. For example, we cannot predict how an animal will respond to the optic flow induced by a banked turn, which contains both translation and rotation self-motion components. To answer this question, it is essential to reveal the underlying neural mechanism, or the algorithm, of optic flow processing. Theoretical studies have described several distinct biologically plausible algorithms ^22–24^. One of these algorithms, the “matched filter” model, suggests that motion receptive fields (RFs) of optic flow sensitive neurons form filters with similar structure to certain types of optic flow, i.e. each RF contains a vector field of locally preferred directions that matches a particular self-motion axis. These matched filters may then form a mathematical basis of self-motion induced optic flow and thus encode translation- or rotation-induced optic flow in a discriminable manner ^22,25^.

In compact insect brains, systematic motion RF characterization has revealed the existence of such RF filters, which match certain types of translational or rotational optic flow ^17,26,27^. However, there are many more neurons involved in optic flow processing in vertebrate brains than in insects. For example, the population of translation- and rotation-sensitive neurons in the pretectal area of larval zebrafish is at least 10 times larger than the population of horizontal system and vertical system neurons in blowfly ^17,18^. It is thus unclear, whether the matched filter algorithm discovered in the compact fly brain is also employed by vertebrate brains. Also, with greater number of neurons comes greater difficulty in systematic RF characterization. Although calcium imaging enables recording of hundreds of neurons simultaneously, no efficient approaches had been available to characterize fine-grained motion RFs in vertebrate brains.

To overcome these limitations, we here made use of contiguous motion noise stimuli and reverse correlation techniques developed recently for the systematic RF characterization of motion-sensitive neurons via calcium imaging ^28^. We measured the fine-grained motion RFs of thousands of neurons in the pretectum and optic tectum of larval zebrafish. The identified RF structures reveal a neural mechanism for robust optic flow encoding in the pretectum. Furthermore, we show that larval zebrafish decompose translation and rotation components in optokinetic and optomotor behaviour as suggested by the underlying encoding mechanisms.

## Results

### Identification of unimodal and bimodal receptive fields sensitive to motion stimuli

To estimate motion receptive fields (RFs), we used a full-surround cylindrical stimulus arena (Figure 1B) and recorded neuronal responses to motion stimuli in zebrafish larvae expressing the calcium indicator GCaMP6s. The stimulus consisted of a contiguous motion noise pattern in which moving directions were spatiotemporally coherent. We then took the average of all stimulus frames that evoked calcium events in a neuron (reverse correlation) and applied bootstrapping statistics to identify which parts of these event-triggered averages (ETAs) can be trusted (Figures 1A, Figure S1 and STAR Methods for details). The method has been described recently ^28^ and enabled us to identify the fine-scale RF structures of 3,926 motion-sensitive neurons from a total of 46,130 region-of-interests (ROIs) recorded in the optic tectum and the pretectal area of seven fish. The shape, span, location, and direction preference of these RFs were highly diverse, as illustrated by the estimated RF structures of the four example neurons in Figure 1D. This method enables inference of preferred motion directions in different parts of the visual field, but is less suitable for estimating the precise diameter of the RF ^28^.

Despite the high structural diversity, we found that most estimated RFs (99 %), contained either one or two significant modes (Figure 1C), i.e. they could be classified into unimodal (or unipartite) and bimodal (bipartite) RFs. The “mode” here is defined as a patch of spatially connected RF components. Although both unimodal and bimodal RFs can be found in optic tectum (OT) and the area pretectalis (APT), most unimodal RFs are located in the OT, and the bimodal RFs are predominantly located in the APT (Figure 1C,E).

### The modes of bimodal RFs are located asymmetrically in the visual field and show patterned direction selectivity

Due to the fundamental structural difference between unimodal and bimodal RFs, we characterized them in different ways. Each unimodal RF had an averaged preferred direction that corresponded to either up, down, left, or right (Figure S2A), and the spatial RF distribution matched results of a previous study ^21^.

For the bimodal RFs, the preferred directions and spatial centres were calculated separately for each mode. To measure the span of bimodal RFs, we calculated the distance between the two mode centres in visual space (intermode distance, illustrated in Figure 2A). The intermode distances ranged from ca. 60° to 315° in our dataset (Figure 2B), and intermode distances of 180° were scarce. The binocular overlap zone was smaller than 30° in our recordings^29^ (cf. illustration in Figure 2C), as the eyes of the whole-mounted fish were in their normal resting position. Therefore, bimodal RFs spanning across two sides of the overlap zone were classified as binocular RFs. Based on this principle and the intermode distance, the bimodal RFs were divided into three groups: 1. Putative monocular RFs receiving visual input from the nasal (−45° to + 45 ° in azimuth) and temporal end (−168° to −90° or 90°to 168°) of a visual hemi-field (shown in red in Figure 2B & 2C); 2. Binocular RFs with short intermode distance (average: ~150°, coloured in green); 3. Binocular RFs with long intermode distance (average ~223°, coloured in blue). For most bimodal RFs, the mode centres were distributed asymmetrically in relation to the anterior-posterior (AP) body axis as shown in Figure 2C.

**Figure 2.**
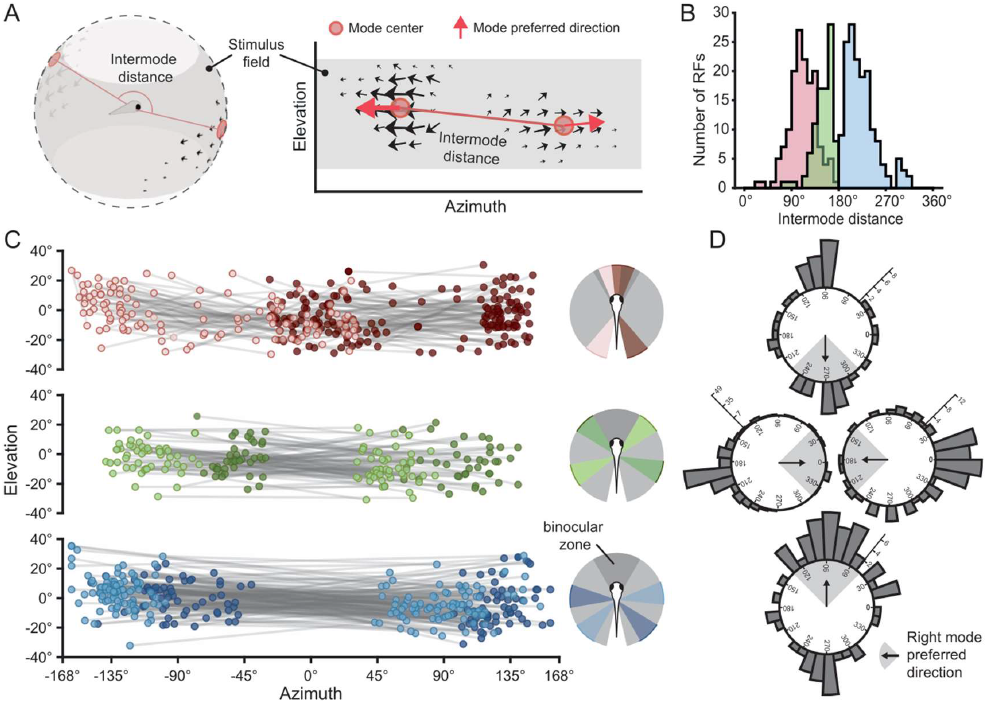
Spatial and directional organisation of bimodal RFs across the visual field. A) Illustration of a bimodal RF projected onto a sphere (left) and the corresponding 2D visual space representation (right). The “intermode distance” was defined as the distance between the two spatially separated components (modes) in the RF. (B-C) Bimodal RFs grouped by their intermode distance and locations in visual field. Each pair of linked dots in (C) indicates the mode centres of a bimodal RF. The RFs tilted to the right hemi-field were coloured darker for clarity. The approximate RF distributions within the visual field are summarized on the right. (D) Pairing of preferred directions (PDs) for all bimodal RFs (i.e. mode pairs). The four polar histograms of the left mode PD are grouped by the PD of the right mode of each RF (indicated by the arrow and shade inside the polar histogram).

Next, we compared the preferred directions within the two modes of each bimodal RF by plotting the distribution of preferred directions in the left mode grouped by their right mode preferred directions (Figure 2D). We found that the RFs preferring leftward (clockwise from above) motion with their right modes are mostly preferring the opposite direction (rightward motion) in their left mode, and vice versa. Among the neurons with vertical direction preferences in the right mode (31 % of all bimodal neurons), 43% of them preferred motion in the opposite direction in the left mode (corresponding to rotation), while the other 57% showed similar direction preference (corresponding to lift translation, Figure 2D).

### Complex bimodal RFs act as linear filters to encode optic flow directions

The directional preferences of the bimodal RFs (Figure 2D) implied these RFs may have similar structures to specific optic flow fields. For example, a bimodal RF that prefers leftward motion in its left mode and rightward motion in its right mode, is similar to the translational optic flow field (TOF) resulting from forward self-motion. And the RFs preferring upward and downward motion in their left and right modes, respectively, are similar to the rotational optic flow field (ROF) resulting from roll. To quantify this similarity to optic flow fields, we calculated the average cosine similarity of all unimodal and bimodal RFs to their closest TOF and ROF (Figure 3A, see STAR Methods section “Fitting optic flow fields to RFs”). The similarity can vary from −1 to 1, where 1 means that the vector field structure of the RF is the same as the corresponding optic flow field, and 0 means the RF is not similar to any optic flow field.

**Figure 3.**
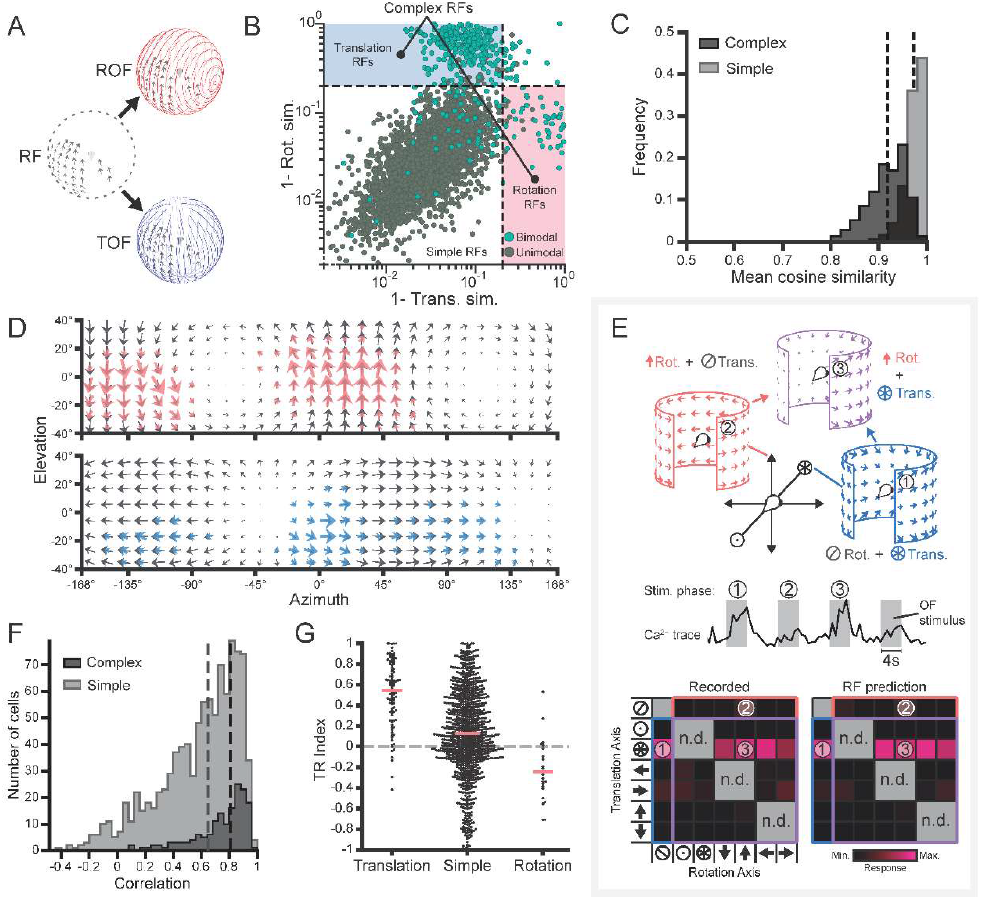
Prediction of optic flow responses in complex RF neurons with the matched filter model. (A) The best matching rotational (red) and translational (blue) optic flow field for an example motion RF (grey). (B) Score of the match of each RF to the fitted rotation (rot.) and translation (trans.) optic flow field. Values below 0.2 (dashed lines) indicate good matched filter RFs; Trans./Rot. sim.: translation or rotation cosine similarity. RFs within blue and red regions were defined as “complex”. (C) Similarities of the complex and simple RFs to the fitted optic flow fields. (D) Two examples of rotation (top) and translation (bottom) RFs (coloured) and the matching optic flow fields (grey); (E) The recorded maximum calcium responses of an example neuron to combinations of different optic flows (OFs, lower left) compared with the matched-filter model predictions (lower right). Three example OFs (1-3) are illustrated at the top and the neuron’s corresponding calcium trace is shown in the middle row; *n.d*.:not determined. (F) The correlation of the recorded and predicted responses for complex and simple RFs. (G) Bee swarm plot of the translation-rotation index (TR index, see also STAR Methods and Figure S2) for neurons with translation, simple or rotation RFs. The dashed lines in C and F and the red horizontal lines in G indicate the median values of the corresponding distributions.

For most RFs (89%), both highly similar TOFs and ROFs existed (cf. dots within the bottom left quadrant in Figure 3B). These RFs were called “simple RFs”, because most of them are unimodal RFs (98%) of relatively simple structure and small spatial extent, and thus they can be interpreted as parts of TOFs and ROFs at the same time. Most of the other RFs were similar to either TOFs or ROFs only (c.f. colour-shaded areas in Figure 3B). These RFs were called “complex RFs” as they are mostly bimodal RFs (319 out of the total 389 complex RFs) containing heterogeneous vector field structures. In addition, most bimodal RFs (80%) are complex RFs. In these complex RFs, the 329 RFs that were more similar to TOFs were called translation RFs while the 60 RFs similar to ROFs were rotation RFs. Two flow field examples for translation and rotation RFs are shown in Figure 3D. Both simple and complex RFs can be well explained by the fitted ROF and/or TOF (Figure 3C): the medians of the cosine similarities for simple and complex RFs are 0.98 and 0.92 respectively.

This finding is consistent with the observation in flies, where the RF organization of direction-selective wide-field neurons were concluded to serve as matched filters indicating the presence of particular self-motions^25^. To test whether or not these complex RFs in larval zebrafish play a similar role, we performed recordings in which 36 distinct optic flow stimuli were presented, followed by the contiguous motion noise (CMN) stimulus (Figure S2C). These recordings lasted 20 minutes. Each optic flow stimulus consisted of either pure translation or rotation, or of a unique combination of translational and rotational optic flow (Figure 3E, also see STAR Methods section “Prediction of optic flow preference”). Using the measured RFs, we applied the matched filter model to predict the calcium responses to the 36 optic flow stimuli for each neuron. The linear correlation of measured and predicted responses to these distinct optic flow stimuli provides a quantitative measure of the prediction accuracy. The average correlation for the neurons with complex RFs (0.81) was higher than the correlation for the neurons with simple RFs (0.65), which means that the complex RFs on average gave better prediction of the neural responses to optic flow stimuli than the simple RFs (Figure 3F and S2D). We quantified the response specificity to translation vs. rotation (TR index, see STAR Methods) and found that bimodal translation neurons had a higher specificity for translation (average TR index >0.5, Figure 3G) than rotation neurons for rotation (TR index ~ - 0.3). The index distribution for simple RFs did not show strong preference to either translation or rotation stimuli as they were also active during presentation of combined rotation/translation optic flow fields.

Together, these results suggest that the identified complex RFs act as matched filters to process optic flow. Most complex RFs prefer either translation or rotation but not a particular combination of the two. Translation neurons were more abundant than rotation neurons and their responses to combinations of optic flow were more predictable with the matched filter model (Figure S2D), suggesting that additional unknown stimulus features modulate the responses of rotation neurons. In the following, we will mainly focus on the population properties of the translation RFs.

### A topographic map of global translation direction in the pretectum

The encoding of global translation direction is essential for any visually guided locomotion. In the sections above, we showed the encoding algorithm (matched filter) implemented by the complex RFs of optic flow sensitive neurons. This allows us to directly infer the preferred translation direction of these neurons as the focus of contraction (FoC) of its best-matching TOF (Figure 4A). Consistent with Figure 2D, most of these neurons prefer translation directions close to the horizontal plane, and another, smaller proportion of neurons prefer vertical translational optic flows (Figure 4A).

**Figure 4.**
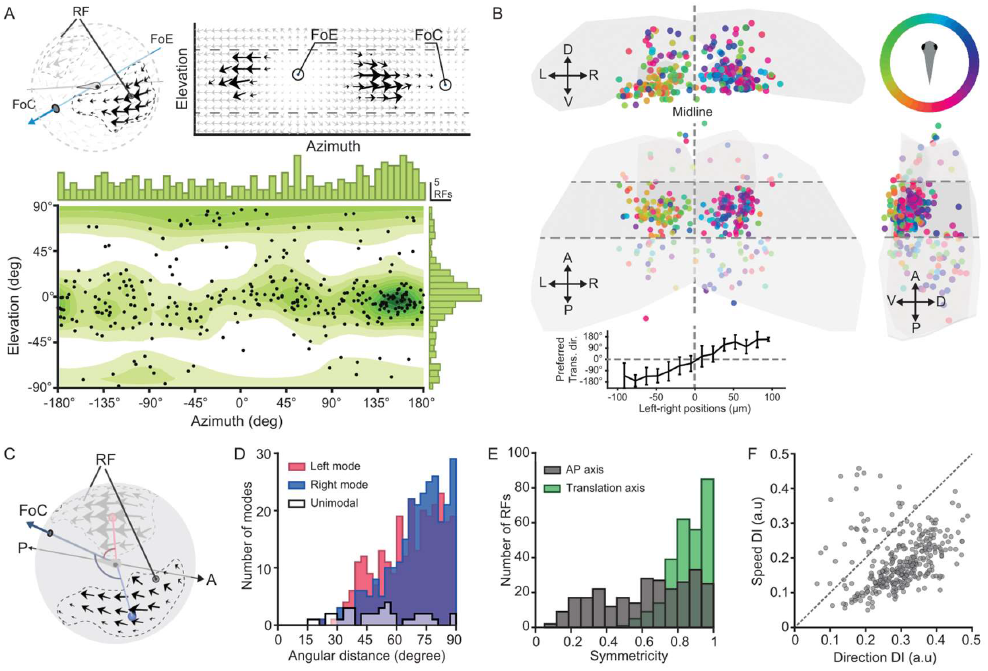
A topographic map of global translation direction in the pretectum. (A) Top: Illustration of focus of expansion (FoE) and contraction (FoC) of the fitted TOF field (grey arrows) of one translational RF (black arrows). Bottom: Preferred translation direction distribution for all translation RFs (n=329). Each dot corresponds to the preferred translation direction (FoC) of a neuron.. For each datapoint, a new decoder was built that used 50 randomly selected RFs (out of the 138 RFs, see STAR Methods) (B) Anatomical locations of neurons (n=329) with translational RFs, coloured according to preferred translation direction in the horizontal plane. The AMC pretectal region is contained within the horizontal dashed lines. The averaged preferred translation directions (trans. dir.) of the neurons binned by the location on the left-right axis (left-right position; bin size = 14.4 μm) are plotted at the bottom, the error bars indicate the standard deviations. See also Video S1 and S2. (C) Illustration of the angular distance between mode centres and preferred translation direction (FoC) analysed in (D). The red and blue arcs indicate the angular distance of left and right mode centres respectively. (D-E). The angular distance of mode centres in relation to the preferred translation direction (D) and the symmetry (E) of mode pairs relative to translation direction and body axis. (F) The discrimination index (DI) for the tuning to translation speed (within the range of 7.5 to 120 °/s) and direction (0-360°) of the 369 translation-sensitive neurons in the AMC of two fish, see also Figure S4 and Video S4.

The majority of these neurons were anatomically arranged in the anterior medial cluster (AMC) region ^18^ of the APT (Figure 4B and S3). These neurons surprisingly formed a topographic map running from left-to-right across the midline according to preferred translation direction, with medial neurons preferring forward translation. This topographic map is structured according to the ego-motion relative to the surround, i.e. a cursotopic arrangement (lat. *cursus* means *running, direction*). Linear regression showed that the preferred translation direction is strongly correlated with anatomical position (R^2^ = 0.73, Figure S4F), and that topography is weaker, but still present, when removing the effects of left-right hemisphere organization (correlation = 0.29, Figure S4G). To visualize this cursotopy further, we recorded the neural responses in the AMC to a TOF stimulus, whose direction changed clockwise (CW) at a constant rate. As expected, the active region in the AMC gradually shifted from left to right, when the translation direction was smoothly changing (Video S1).

There are different ways in which a translation-sensitive neuron could detect the direction of translational optic flow. For example, its complex RF could sample the vicinity of the FoC and/or FoE, or the regions away from FoC and FoE, which contain stronger and more coherent local motion. To identify how the complex translation-sensitive RFs solve this task, we calculated the angular distance between RF mode centres(s) and the preferred translation directions, including both unimodal and bimodal RFs in the analysis. We found that the mode centres of most bimodal translation RFs were located 60-90 degrees away from the preferred translation direction (Figure 4D), corresponding to regions with highest speed and strongest local motion coherence in the optic flow field. Furthermore, we found the mode centres of most bimodal translation RFs, which distributed asymmetrically in relation to the body axis (cf. Figure 2C), to distribute symmetrically in relation to the preferred translation directions (Figure 4E). Unimodal complex neurons, however, were distributed more widely and located slightly closer to the FoC or FoE (~51° distance on average).

The responses of translation-sensitive neurons in the AMC were modulated not only by translation direction, but also speed. Neurons were jointly tuned to translation direction and speed (Figure S4) and preferred speeds from approximately 15°/s to 40°/s. To estimate the fidelity of the joint speed and direction representation in the AMC neurons, we computed the discrimination index for translation direction and speed for each neuron. A higher discrimination index means the neural responses are more discriminable, which in turn, indicates the neuron is more strongly tuned to the variable. We found that almost all neurons were tuned to both translation direction and speed, while the tuning discriminability for translation direction was slightly higher than that for translation speed in most neurons (Figure 4F).

### Robust encoding of translation direction

As shown above, neuronal responses to optic flow can be faithfully predicted by the matched filter algorithm based on the measured pretectal RFs (Figure 3). An essential question to ask is whether the information encoded by neurons with translation RFs is sufficient to estimate the translation direction when distractors or noise are present? In other words, can an animal robustly estimate its ego-motion based on the neural responses of these translation-sensitive neurons in the AMC region?

To answer this question, we first constructed a population decoder to quantitatively determine whether the encoding of translation direction is robust against rotational self-motion interference (see STAR Methods section “Population decoder for self-motion estimation”). Translational optic flow fields are greatly affected by additional rotational optic flow components. For example, added rotation will cause an apparent shift of the FoC point and changes of spatial symmetry. This decoder was a variant of the maximum-a-posteriori decoder that used the measured RFs (i.e. using data from Figure 3E) to predict responses to different TOFs. The decoder then estimated the translation direction of the presented optic flow combination by minimizing the squared difference between the recorded responses of the translation-sensitive neurons and the prediction by their RFs (Figure 5A, see STAR Methods for details).

**Figure 5.**
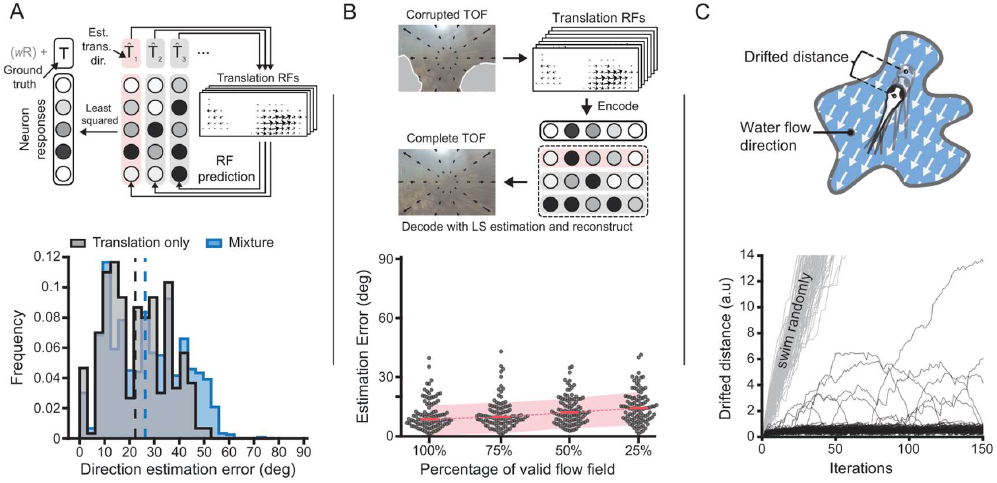
Robust encoding of translational optic flow direction by translation RFs. (A) Top: Illustration of the constructed population decoder estimating the stimulus parameters from the responses of optic flow sensitive neurons (see also STAR Methods and Figure S5). The “w” and “R” in the top left indicate the rotation weight in relation to translation and rotation parameters respectively; Est. trans. dir.: estimated translation direction; Bottom: The decoder estimated the translation direction for translational optic flow with (“mixture”) and without (“Translation only”) added rotational components. A histogram of estimation errors is shown as well as the median errors (dashed lines). For each datapoint, a new decoder was built that used 50 randomly selected RFs (out of the 138 RFs, see STAR Methods). (B) Top: Illustration of the autoencoder constructed with the translation RFs for determining the direction of corrupted translational optic flow fields (TOFs, see STAR Methods). Bottom: bee swarm plots showing the estimation errors for the TOFs with different degrees of visual field corruption. The dashed line and shaded area indicate the median ± the standard deviation. See also Figure S5. (C) The drifted distance over time of 100 simulated fish (black lines) using the population decoder in (B) to estimate and swim against the direction of ego motion (illustrated in the upper drawing). The grey lines correspond to the drifted distance when the simulated fish swam randomly. See also Video S3.

To our surprise, the estimation error for the translation direction was similar with or without rotational optic flow interference, which suggests a robust encoding of translation direction (Figure 5A bottom). Indeed, AMC responses to the pure translational optic flow and the mixed optic flow looked very similar (Video S1 and S2). In other words, the translation information can be extracted from the mixed optic flow field solely with the information encoded by the translation RFs in the AMC region.

To compare the encoding quality of the speed and direction information, we constructed a separate population decoder (Figure S5A-B, see STAR Methods for details). In contrast to the low decoding error for translation direction (median error: 6.6 degrees, the decoding error for translation speed was high (8.56 degrees/second or 32.6% of the stimulus speed) when compared to the range of encoded speeds.

In the natural environment, optic flow is often corrupted by occlusion, the lack of visual cues or by interfering object motion. Since it is essential for animals to determine their moving direction from corrupted optic flow, we determined to what extent this is possible using translation RFs. For this purpose, we constructed an autoencoder whose encoder and decoder employ the same response functions determined by the corresponding translation RFs (Figure 5B top).

We first tested the encoding quality for translation optic flow fields (TOFs) with 100 different translation directions at 4 completeness levels (25 %, 50 %, 75 %, 100 %), corresponding to the fractional area of the visual field in which unblocked optic flow is available. For example, 25% indicated the optic flow in 25% of the visual field is visible. For each TOF, an autoencoder was constructed with 50 RFs randomly sampled from the total 329 translation RFs (see STAR Methods for details). As expected, the estimation error gradually increased with the decreasing completeness levels, however even for the 25% completeness level in which most of the optic flow field is missing, the median estimation error was still below 20 degrees. It was only 5.7 degrees higher than the one for the 100% completeness level (8.7 degrees, Figure 5B bottom), suggesting that the corresponding RFs are well prepared to encode self-motion direction in environments with sparse motion information. We also tested the encoding robustness against moving objects, which corrupt optic flow fields with wrong motion information. For this type of corruption, the decoding was less robust, especially when half or more than half of the visual field was covered by moving objects (Figure S5E). This poor performance was not surprising, since it is hard to distinguish the cues for global motion and object motion purely from the optic flow fields without additional information.

The robust encoding of translation direction is critical for many locomotor behaviours, e.g. for self-stabilizing behaviour in water currents ^30^. The continuous change of the water current direction and the relative distance to the surround alter the optic flow continuously, which requires fish to constantly adjust their swimming direction to maintain their body position. We were curious to see, whether these translation RFs are capable of such a complicated task. We therefore constructed 100 randomly shaped “ponds”. In each of these simulated ponds, the water flowed at constant speed (1 a.u./iteration) in a random direction. The simulated fish then swam at the same speed into the directions estimated by the autoencoder from the instantaneous optic flow fields (Figure 5C, also see example fish in Video S3). In most simulations, the total drifted distances were smaller than 1 a.u., which suggested that the translation RFs accurately encoded sufficient information in this highly dynamic process to determine appropriate translation directions for self-stabilization behaviour.

### Distinct optokinetic and optomotor responses to translational and rotational components in complex optic flow fields

Earlier, we asserted that without systematic characterization of neural mechanisms, we cannot even answer how fish will respond to the optic flow during a bank turn as it consists of both translation and rotation (Figure 6A). The revealed matched filter algorithm provides insight in the self-motion extraction from optic flow in zebrafish larvae. The robust encoding of translation direction (Figure 5A and Video S2) against the interference of rotational optic flow component strongly suggests translation and rotation information from the optic flow input are segregated in the zebrafish brain. It is thus very likely that fish will respond separately and simultaneously to the translation and rotation related optic flow components during any movements involving both translation and rotation.

**Figure 6.**
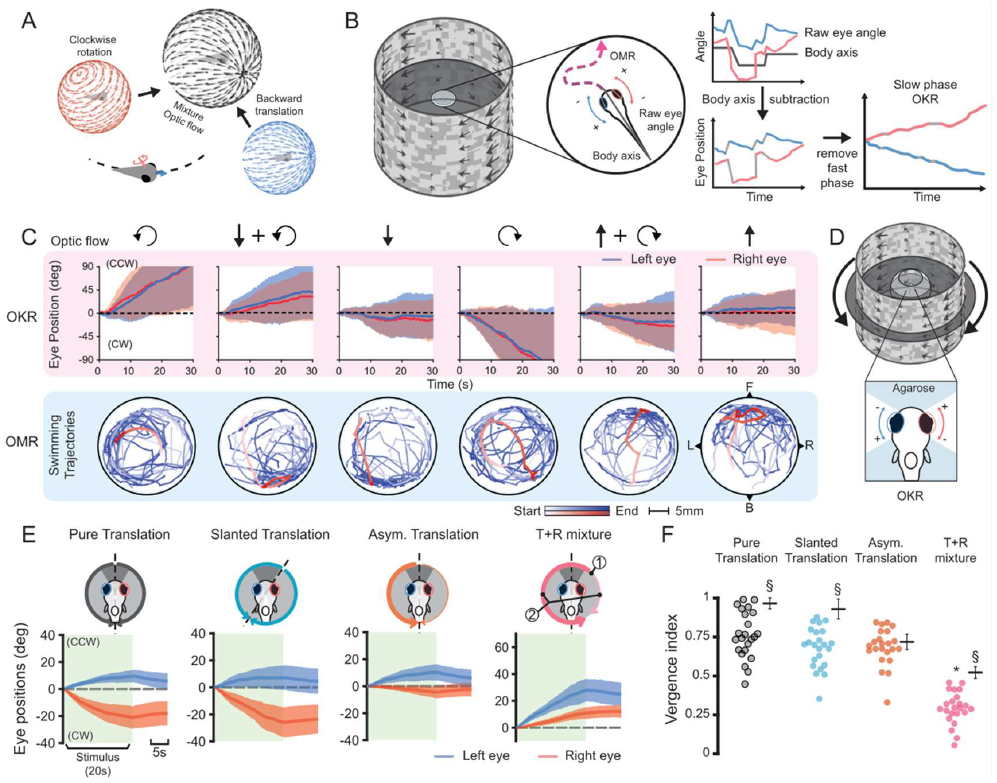
Zebrafish decompose mixed rotation and translation optic flow in optokinetic and optomotor responses (OKR, OMR). (A) Illustration of the mixture optic flow field resulting from the counter-clockwise (CCW) turning of a forward moving fish. (B) Setup for simultaneous OKR (eye movements) and OMR (swimming trajectory) recording for freely swimming fish (see also STAR Methods and Figure S6). For OKR (right), saccades were removed to analyse cumulative slow phase eye movements. (C) The OKR and OMR upon optic flow stimulation containing translation and/or rotation components in 24 fish (black arrows on top indicate the OF stimulus composition; solid line: mean; shaded area: standard deviation across all fish). One example swimming trajectory is coloured in red for each stimulus. “F”, “B”, “L”, “R”: the forward / backward / left / right direction, respectively. (D) OKR recording setup for agarose-mounted “semi-restrained” fish in a spherical water container. The agarose around the eyes was removed to allow free eye movement. (E) The eye position traces (mean ± standard deviation) across 18 trials recorded in five semi-restrained fish. The OF stimuli are illustrated on top. Dark grey corresponds to binocular overlap region and the curvy arrows in D and E indicate the average motion in each hemi visual field. Arrow thickness corresponds to the relative stimulus speed. ① and ② label two local features present when adding rotation on top of translation OF: the focus of expansion (FoE) shifts and speed in the two visual hemi-fields is unequal. (F) Ocular vergence index for different OF patterns (based on eye traces in (E)). For comparison, we also simulated the vergence index (black mean ± std. bars) expected for independently moving eyes responding to the average stimulus directions and speeds in each hemi-field (using bootstrapping of local stimulus statistics, see STAR Methods). 1 and 0 correspond to pure convergent/divergent and pure conjugate movement, respectively. The asterisk indicates significantly lower vergence indices compared to the other three groups. The “§” indicates a significantly lower measured ocular vergence index in comparison to the corresponding local flow statistic. The Ranksum test (α= 0.05) and Bonferroni correction was used to assess significant differences.

To test this, we selected the optomotor responses (OMR) and optokinetic responses (OKR) as two behaviour readouts as they are tightly linked to optic flow perception and represent prominent stabilization behaviours in zebrafish ^18,30^. We recorded the OKR and OMR of 24 freely swimming fish to pure forward/backward translational and CW/CCW rotational optic flow, as well as to combinations of both (Figure 6B-C).

If fish failed to decompose translational and rotational components, we would expect inappropriate behavioural responses, for example the occurrence of rotational behaviour to translational optic flow. In our experiments, fish responded to pure rotation with both OMR and OKR, as indicated by their circular swimming trajectories and conjugate eye movements (Figure 6C and S6). For the pure translation optic flow, fish swam with the optic flow direction, while clear conjugate or vergence eye movements were absent.

If fish were unable to extract both translational and rotational self-information from the combinatory optic flow, their responses should be similar to the responses evoked by pure translational or rotational optic flow. Interestingly, when combinations of translation and rotation stimuli were presented, fish clearly swam with the translation direction while performing conjugate eye movements in the rotation direction, indicating the perception of both translation and rotation components in the optic flow stimuli (Figure 6C and S6).

Since OKR during translation stimuli may be disrupted by changes of head direction in freely swimming fish, we repeated the experiment in semi-restrained fish, which could not swim but were able to move their eyes (Figure 6D). In contrast to the results in freely swimming fish, the semi-restrained fish performed clear vergence eye movements to pure translational optic flows. For optic flow combinations, both semi-restrained and freely swimming moving fish showed conjugate eye movements without significant vergence components (Figure 6E). Together, this behavioural evidence suggests that translation and rotation components are perceived separately and simultaneously by zebrafish.

There are at least two local features that are different between the pure translational and the mixed optic flow: the FoC/FoE position and the average motion speed of the left and right visual hemi-fields (cf. upper right illustration in Figure 6E). The head direction invariant OKR implied that the distinct processing of translation optic flow with or without rotational components does not rely on particular local motion features. To test this, we constructed two variants of translational optic flow. In the “slanted translation” variant, the translation direction of the pure translational optic flow field was shifted, so that its FoC was at the same position as the one in the mixture of translation- and rotation-induced optic flow. The average speed in the left and right halves of the “asymmetric translation” optic flow fields was altered to mimic the different speed in the mixture optic flow field. The OKRs to these stimuli were slightly different from the ones to pure translation optic flow, however, none of them could be classified as conjugate eye movements (Figure 6E).

We quantified the extent of conjugate vs. vergence eye movements using a vergence index (Figure 6F, 1 = pure vergence movement; 0 = pure conjugate movement; see STAR Methods for details). The vergence index for the translation optic flow combined with a rotation component was significantly shifted towards conjugate eye movements in comparison to all other translation stimuli, while the vergence indices for the various translation stimuli were similar to each other (Figure 6F, Wilcoxon rank-sum tests). Furthermore, each vergence index (except for the “asymmetric translation” stimulus) was slightly shifted towards conjugacy in comparison to what would be expected based on the distribution of local velocity vectors in the optic flow field (see black crosses in Figure 6F which show the bootstrapped null distribution). Together, these results show that the animal cannot be tricked into processing rotational motion when only certain local aspects of rotational flow are implemented in overall translational flow patterns.

## Discussion

Via the systematic characterization of the fine-grained receptive field structure of motion-sensitive neurons in the optic tectum and pretectal area, we revealed how self-motion information can be extracted from the complex RFs, and how the encoding properties may be explained by the matched filter model. We furthermore demonstrated the robust encoding of self-translation direction and speed in the translation-sensitive neurons in the pretectal AMC region. Finally, our behavioural results show that zebrafish optokinetic and optomotor responses draw on the decomposition of rotational and translational information.

### Complex RFs implement the optic flow detection algorithm

To our knowledge, this is the first study that systematically estimated the detailed RF structure of optic flow processing neurons in a vertebrate animal model. The feasibility to estimate the 3,926 motion RFs reported here, hinged on the availability of a fast, multiplexed RF estimation technique developed for calcium imaging ^28^. The RF structures of the global motion sensitive neurons revealed in this study showed remarkable similarity to translational and rotational optic flow fields (Figure 3C). This observation is consistent with the “matched filter” model proposed in flies, which suggests optic flow sensitive neurons extract self-motion information by matching their RFs with impinging optic flow fields ^17,25,31^. The calcium responses of our complex RF neurons, especially the ones with translation RFs, are well described by this model (Figure 3F, Figure S2D) and the level of remaining unexplained response modulation is small for most of the complex RF neurons. Most complex RFs are bimodal RFs (Figure 3B) and it is possible that the bimodal RFs of these neurons are inherited from pairs of presynaptic unimodal RF neurons^32^, which may be confirmed by systematic connectivity characterization in the future.

In contrast, the responses of the simple RF neurons frequently differed from the predictions of the matched-filter model (Figure 3F, Figure S2D), and the precise choice of thresholds in our analysis did not appear to have a major effect on the matched-filter properties of neurons (data not shown). One possibility is that many of these “simple” neurons had non-linear RF structure (i.e. suppressive components) remaining undetected in our RF estimation. Indeed, previous studies reported that about 80% of the local motion-sensitive neurons in zebrafish optic tectum and salamander retinal ganglion cells (RGC) were suppressed by wide-field motion ^21,33^. An alternative (and not mutually exclusive) explanation for the observed prediction discrepancy is that our simple RF neurons carry out functions in the visual system, which are only distantly related to optic flow encoding. Further investigation is needed to characterize inhibitory components and functions of simple RF neurons in the brain.

### A topographic map of preferred translation directions in the AMC pretectal region

Within the pretectal AMC region, translation-selective neurons are roughly arranged according to preferred translation directions in a single topographic map that spans both pretectal hemispheres (Figure 4B & S4F-G, Video S1 & S2). Previous reports already revealed differential directional processing across pretectal locations and related these direction-selective neurons to the triggering of optomotor responses ^18-21,34^.

These results agree with our findings despite major differences in employed visual stimulus setups. In these studies^19,34^, motion stimuli were projected on a flat screen from below, which can be interpreted as the bottom part of a translational optic flow field. For example, the nasal-temporal-ward motion covering the bottom visual field can be interpreted as the bottom part of a translational optic flow field resulting from forward self-motion. Using this interpretation, the topographic map of translation direction found in this study (Figure 4B) is consistent with the direction map described in the previous study^19^ (Ibid. Figure 3a) in terms of the preferred direction in different regions of the AMC. Furthermore, our results show that this pretectal structure encodes the translation axes of optic flow rather than local motion directions for each eye. Unfortunately, RF structures in the peripheral lower visual field (<-40° elevation) could not be estimated due to the hardware limitation in our study. A whole field stimulus display setup will be desired for estimating the complete RF structure in the entire visual field.

Besides translation direction, speed is another critical factor for locomotor behaviour. Larval zebrafish can adjust their swimming behaviour to match the speed of wide-field visual motion stimuli ^35,36^. Here we show that most translation-sensitive neurons in the AMC region are jointly tuned to both translation direction and speed (Figure 4F). The speed information in translational optic flow is usually conjugated with the depth and location of stimuli in the visual field ^23^. Whether and how translation-sensitive neurons in the AMC regions untangle this information for determining self-motion speed requires further investigation.

### Represented optic flow axes

We noticed some invariant properties across RFs in the optic tectum and pretectum: I) the preferred directions for both the unimodal and bimodal RFs are biased toward either horizontal or vertical motion, with only a few neurons responding to oblique motion in Figures 2D and S2A ^20,37,38^; II) bimodal vertical neurons together encode translation and rotation, whereas most bimodal horizontal neurons encode translation (Figure 2D); III) the modes of bimodal translation RFs are mostly distributed symmetrically with respect to the preferred translation direction instead of the body axis (Figure 2C and 4E); and IV) most translation RFs sample regions to the side of the preferred translation direction, where motion coherence is high (Figure 4D).

It is unclear at this point, what the specific advantages of this layout are. The layout results in a relatively even distribution of neuronal preferred directions in the horizontal plane, while translation neurons preferring oblique translation directions outside the horizontal plane or the vertical axis are absent. One possibility is that this representation serves to improve the coding efficiency by balancing the needed number of active neurons (sparseness) and the needed total neuron number: the orthogonal representations could form a mathematical basis, so that oblique directions could be encoded by the activity of (at least) two neurons with orthogonal preferred directions. Our decoder simulations show that the relatively rich representation of horizontal directions can be beneficial for targeted locomotor responses in the horizontal plane, e.g. during OMR (Figure 5C). However, spontaneous swimming as well as prey capture behaviour includes frequent locomotor deviations from the horizontal plane ^39,40^ as well. The surprising scarcity of representations of yaw rotation suggests that it is negligible in visual brain areas of the zebrafish. Possibly, inhibitory outputs of translation-selective neurons or the activity in the vestibular system could be integrated in downstream neurons to signal the presence of yaw rotation. Furthermore, each eye can, to some extent, correct for horizontal image slip by itself, potentially making a binocular rotation representation dispensable.

### Decomposition of translation and rotation self-motion components

While evidence for separate processing of translation and rotation information in the corresponding neural correlate has been presented before ^12,18,20,41^, our study shows that translation/rotation sensitive neurons are not exclusively responsive to pure translational/rotational optic flow, but also respond to the mixed optic flows in a predictable manner (Figure 3). Furthermore, the population encoding of translation direction is robust against rotation interference (Figure 5A and Video S2), which indicates that the zebrafish pretectum separates translational and rotational self-motion components and is biased towards the identification of translation components. In contrast to translation RFs, the responses of rotation-sensitive neurons to mixed optic flow stimuli are strongly modulated by the non-preferred (translation) components and poorly explained by their RF structure (Figure S2D). It remains to be answered whether these rotation-sensitive neurons in zebrafish are similar to the neurons jointly representing translation and rotation components in primate parietal lobes^12^, and what role they do play in optic flow processing.

The decomposition of optic flow requires the translational and rotational self-motion information not only to be separated, but also to be utilized differentially in behaviour. Our behavioural experiments revealed evidence of this decomposition, since fish swam toward the corresponding translation direction implemented in the optic flow field (OMR, Figure 6C) while spinning both eyes in the direction of rotation (OKR, Figure 6C, E and F). The OKR is more sensitive to the rotation component of a combined optic flow stimulus (Figure 6F), suggesting that OKR and OMR might show performance differences for different types of optic flow. Our experiments on isolated local features (Figure 6E&F) suggest that the integration of local features but not a specific local feature is essential for the optic flow processing in zebrafish.

We hypothesize that our identified population of pretectal neurons form a visual encoding pivot, where complex RFs directly extract and convey the translation direction to the premotor areas. It will be interesting to determine the full neural circuit for this optic flow decomposition in the future, including the connectivity of translation-sensitive neurons in AMC ^32^

In summary, we have systematically characterized the receptive field properties in zebrafish visual brain areas that help us to reveal a neural mechanism for optic flow processing. Our results suggest that individual neurons form matched filters to extract translation and rotation self-motion components from optic flow. We identified an anatomically ordered arrangement of different functional cell types, which, as a population, decompose rotational and translational self-motion components of the animal. Future work is needed to show, how these cellular building blocks are wired together in a hierarchical circuit to mediate visually driven reflexes and voluntary behaviours.

## Acknowledgments

We thank Thomas Nieß (glassblower shop, University of Tübingen) and Klaus Vollmer (fine mechanics workshop, University of Tübingen) for technical support, and Herwig Baier (MPI of Neurobiology) for providing the Tg(elavl3:nls-GCaMP6s)mpn400 zebrafish line. We thank Holger Krapp (Imperial College London), Eva Naumann (Duke School of Medicine), Julian Hinz (Friedrich Miescher Institute for Biomedical Research) and members of the laboratory for feedback on the manuscript.

This work was funded by the Deutsche Forschungsgemeinschaft (DFG) grants EXC307 (CIN - Werner Reichardt Centre for Integrative Neuroscience) and INST 37/967-1 FUGG, and a Human Frontier Science Program (HFSP) Young Investigator Grant RGY0079.

## Author contributions

Conceptualization: Y.Z. and A.B.A., Methodology: Y.Z., R.H., A.B.A, Software: Y.Z., R.H., Formal Analysis: Y.Z., A.B.A., Investigation: Y.Z., R.H., W.N., Writing - Original Draft: Y.Z., A.B.A., Writing - Review& Editing: Y.Z., R.H., W.N., A.B.A., Visualization: Y.Z., Supervision: A.B.A., Y.Z., Funding Acquisition: A.B.A..

## STAR Methods

### KEY RESOURCE TABLE

| REAGENT or RESOURCE | SOURCE | IDENTIFIER |
| --- | --- | --- |
| Chemicals, peptides, and recombinant proteins |  |  |
| 10000 MW Texas Red dextran | Thermo Fisher | https://www.thermofisher.com/ |
| Deposited data |  |  |
| Max Planck Zebrafish Brain Atlas | [47] | https://fishatlas.neuro.mpg.de |
| Processed data for generating the figures contained in this manuscript | This paper; G-Node GIN | https://doi.org/10.12751/g-node.hkzu3l |
| Experimental models: Organisms/strains |  |  |
| Tg(*elavl3:nls-GCaMP6s*)*mpn400* | [48] | ZFIN ID: ZDB-ALT-170731-37 |
| *mitfa* | [49] | ZFIN ID: ZDB-GENE-990910-11 |
| Software and algorithms |  |  |
| MATLAB (data analysis) | MathWorks | https://www.mathworks.com/products/matlab.html |
| MOM® Computer System (MCS) | Sutter instruments | https://www.sutter.com/MICROSCOPES/mcs.html |
| Code for generating the figures contained in this manuscript | This paper; G-Node GIN | https://doi.org/10.12751/g-node.hkzu3l |

### RESOURCE AVAILABILITY

#### Lead contact

Further information and requests for resources and reagents should be directed to and will be fulfilled by the lead contact, Aristides B. Arrenberg.

#### Materials Availability

This study did not generate new unique reagents

#### Data and code availability

The scripts and the required pre-processed dataset for the figures and supplementary figures included in the current study are available in a public repository (https://doi.org/10.12751/g-node.hkzu3l)

The raw calcium imaging data and animal behaviour videos supporting the current study have not been deposited in a public repository because of their large size but are available from the corresponding author on request.

### EXPERIMENTAL MODEL AND SUBJECT DETAILS

#### Animals

Zebrafish (Danio rerio) were maintained on a 14/10h light-dark cycle at 28°C. The transgenic zebrafish line *Tg(elavl3:nls-GCaMP6s)mpn400* was used for the *in vivo* two-photon calcium imaging (Dal Maschio et al. [45]). The top 10 - 20% of larval zebrafish with high calcium indicator expression levels and strong optokinetic responses (OKR) to moving gratings were selected from all hatched fish at 4-day post-fertilization (dpf) for *in vivo* calcium imaging experiments.

5-7 dpf zebrafish larvae carrying mutations in the *mitfa* gene (nacre) were used for OKR and optomotor responses (OMR) recordings. Again, the top 10-20% larval zebrafish sensitive to visual motion stimuli indicated by their strong OKR to moving gratings were selected at 4 dpf for these behavioural experiments.

All behavioural and *in vivo* calcium imaging experiments were performed on 5-6 dpf larval zebrafish at room temperature.

All animal experiments were licensed by the local authorities (Regierungspräsidium Tübingen) in accordance with German federal law and Baden-Württemberg state law.

## METHODS DETAILS

### Visual stimulation setup

All visual stimuli used in this study were displayed on a custom-built cylindrical green LED arena (160 mm height, inner diameter = 198 mm) covering 336°-by-80° of the visual field (−168° to 168° in azimuth; −40° to 40° in elevation). The 2 halves of LED arena consisted of 224 square LED tiles (8 rows × 14 columns × 2 arena halves); each tile had 8×8 evenly distributed LEDs emitting at 570nm (Kingbright TA08-81CGKWA). Each LED covered ~1.5 visual degrees on average, note the coverage on vertical direction varied slightly with the latitude of LEDs. All stimuli displayed in the arena had 100% contrast as the LEDs in the stimuli were either on or off. All LED tiles were covered with a diffusion filter foil (LEE no. 252, article 595-1780-2520) and a high-pass filter foil (LEE no. 779, article 595-1700-7790, castinfo.de, Hagen, Germany) for light homogenization and light interference reduction in calcium imaging respectively.

The custom software developed in previous studies (Joesch et al. [46]; Reiser and Dickinson [47]) was used for stimulus uploading and display control. To minimize the interference of LED light with the calcium imaging, the LEDs were flickering in a duty cycle (~1 ms) to be switched on only during the fly-back time (approx. 160 μs) of the scanning mirrors in-between line scans.

### Contiguous motion noise stimulus

The contiguous motion noise (CMN) stimulus for motion RF estimation was constructed as described in our previous study ^28^. Briefly, the optic flow in the CMN stimulus was determined by a three-dimensional (8 rows × 28 columns x 12000 frames) motion vector noise matrix, in which motion vectors were spatiotemporally correlated. The vector length and direction indicate the motion speed and direction in the corresponding local region. The spatiotemporal correlation in the noise was achieved by convolving a white vector noise matrix with a multidimensional Gaussian kernel (μ_x,y,t_ = 0, σ_x, y_ = 18.78 visual degrees, σ_t_ = 0.33 seconds). The spatiotemporal correlation in our stimulus may introduce bias to the RF estimation, i.e. overestimation of the local coherence in the RF. The spatiotemporal extent of this bias was quantified as the contiguous radius (CR) of the CMN stimulus for which the pairwise vector correlation surpassed 0.1. The CR is calculated as:

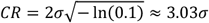

The CR in spatial and temporal domain are therefore 57 (3.03 × 18.78) visual degrees and 1 (3.03 × 0.33) second respectively.

The noise vector matrix was then transformed to a 64 (row) × 224 (column) × 12000 (frame) binary movie displaying at 15 frame-per-second (fps) with the Reichardt encoder described previously (Zhang & Arrenberg, 2019). Each motion vector in the noise vector matrix was approximated by the displacement of the binary noise pattern displayed on an 8×8 LED patch at the corresponding position on the arena. The spatial frequency of the 224 binary noise patterns (one for each LED patch) was set to 0.1 cycles per visual degree which is optimized for visual motion perception in larval zebrafish ^48^.

The speed distribution of the CMN stimulus follows the Rayleigh distribution:

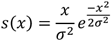

where σ = 12.35 visual degrees/second. Therefore, the speed in 95% of the pixels in our CMN stimulus movie was contained in the range between 0 to 30 visual degree/second.

### Optic flow stimuli design

Similar to the CMN stimulus, the optic flow stimuli were designed in vector field forms with the same spatial resolution (8×28) as in the CMN stimulus, and they were converted to binary noise movies by the same Reichardt encoder to be displayed on the LED arena. The rotational 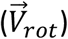 and translational optic flow vector fields 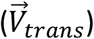 were constructed as following:

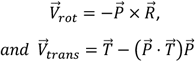

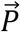 is a unit vector point from the observer (the origin) to the location *P* in the visual field. 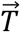 and 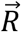 are the unit vectors indicating the translation direction and the rotation axis of the translational and the rotational optic flow fields respectively. The operations <x> and <·> are the cross and dot products of two vectors respectively. And a mixture optic flow containing both translation and rotation components can be obtained as:

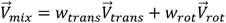

where the *w_trans_* and *w_rot_* represent the relative strength of the translation and rotation components in the mixed flow field.

The speed of motion cues in a translational optic flow field depends on both their locations in the optic flow field and the distance to the observer. As this study is not focused on the distance effect, all motion cues in our optic flow fields were set to be equally distant from the observer. To correct the projection distortion of the optic flow fields on the cylindrical LED arena, the optic flow fields calculated above 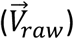 were transformed as following:

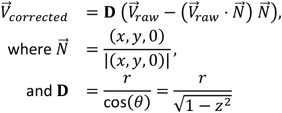

where the *x, y, z* are the LED locations in the Cartesian coordinate system where the origin is the location of the observer, and θ is the corresponding elevation angle.

In the translational optic flow fields, the motion speed is maximum at the locations S which are perpendicular to the translation direction 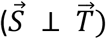. For convenience, the translation speed of a translational optic flow field was defined as the speed in these locations (colour-shaded circles in the illustrations of Figure S4A).

### *In vivo* two-photon calcium imaging

The selected fish were immobilized in 1.7% low melting-temperature agarose (wt/vol, E3 medium) at the tip of a transparent plastic triangle stage (tip angle < 45%, Figure S2E) on the experiment day. The triangle stage with the mounted fish was later placed in the centres of a spherical glass container filled with transparent E3 medium. Then the water container was fixed on a metal holder placed in the centres of the LED arena. The body position and orientation of the mounted fish was adjusted so it was centred in the LED arena with the dorsal side facing up (roll correction) and the nose pointing straight to the front (0° azimuth and elevation) without any tilting.

The calcium activities in neuron somata were recorded with a movable objective microscope two-photon setup (Sutter Instruments, Novato, California, USA; Euler et al. [49]) coupled to a Coherent Vision-S Ti-Sa laser. The emitted fluorescence light of the calcium indicator protein, GCaMP6s, triggered by the excitation laser beam at 920 nm (prepulse compensation: 9756 fs^2^, laser power = ~33 mW at object plane) was collected with a 20x/1.0 Zeiss objective using an emission dichroic with bandpass 500-550 nm. The image time series were recorded using MScan software (Sutter instruments). The calcium images acquisition rate was 2 fps at 512 × 512 pixel^2^ and a magnification of 2 (0.44 × 0.44 μm^2^ per pixel).

### Recording behaviour in freely moving animals

The fish at 5-6 dpf with strong behavioural responses to moving gratings were selected on the days of OKR or OMR experiments. To determine the optomotor responses of freely swimming larvae to optic flow stimuli, we built a cylindrical swimming reservoir (diameter: 20mm, depth: 3mm) on the centres of a 50mm-diameter round transparent plastic plate. To make the stimulus displayed on the LED arena completely visible, the cylindrical reservoir wall was made from 2% low melting-temperature agarose (i.e. no plastic wall) to prevent plastic-agarose interfaces, which would introduce additional optical artifacts. This reservoir was placed in the centres of the LED arena, whose radius was >9 times larger than the radius of the reservoir. Because the optic flow stimuli were all symmetric to the horizontal plane, we do not expect any strong disturbance from internal reflection to animal behaviour as discussed in a previous paper ^50^. A selected fish was then placed in the reservoir filled with E3 medium.

An infrared LED lamp (850 nm, Conrad, Item No. 491248-62) was placed above the LED arena to provide relative homogeneous light for illumination. A CMOS camera (DMK23UV024, The Imaging Source GmbH, Bremen, Germany) with a 6 mm lens (C-Mount Lens FL-HC0614-2M, Ricoh) was placed at 30 mm below the reservoir. Fish behaviour was recorded in 100 frame-per-second (fps) with a custom written Python application.

The body axis, body position, and raw eye angles were extracted offline by a custom written OpenCV based Python script. Briefly, the eye pixels and body pixels were extracted using a threshold-based segmentation method. The eye and body centres positions were computed as the first moment of the corresponding pixel ensembles. To obtain the body axis or the head direction, we first applied the principal component analysis (PCA) to the body pixel ensemble to obtain the rough body orientation as the first principal component (PC) vector. The coarse body direction (the sign of the first PC vector) was then determined as the side with higher variation on the second PC axis. We further refined the body axis by computing the vector that is perpendicular to the line linking the centres of two eyes and pointing into the coarse body direction determined previously. The orientation of the two eyes was also computed with PCA in the same manner. The eye position was computed as the angular difference between the eye orientation and the body axis (Figure 6B). The eye traces were first smoothed with a moving mean filter with the window size of 25 frames (250 ms) to reduce noise interference. In order to extract the OKR-related slow phase eye movement, the eye angle changes faster than 90 degrees/second were set to zero. The total eye angle changes over the entire stimulus period were computed for the left and right eyes (Δleft and Δright).

### Eye movement recording in mounted fish

The selected 5-6 dpf fish were immobilized in 1.7% low melting-temperature agarose (wt/vol, E3 medium) at the tip of a transparent plastic triangle stage (tip angle < 45%) on the experiment day. The agarose surrounding the eyes of the mounted fish were carefully removed to allow free eye movement. The triangle stage with the mounted fish was later transferred into a spherical glass container filled with E3 medium. The container was placed in the centres of the cylindrical LED arena. The body position and orientation of the mounted fish was adjusted later, so it was in the centres of the LED arena with the dorsal side facing up (roll correction) and the head pointing to 0°elevation (pitch correction) and 0° azimuth (yaw correction).

A CMOS camera (DMK23UV024, The Imaging Source GmbH, Bremen, Germany) was placed at ~7 centimetres below the mounted fish (~3 cm below the bottom of the glass bulb) to record the horizontal eye movement. To create homogeneous light for eye illumination, a 3-cm-width white paper stripe was placed above the opening of the glass bulb to diffuse the light emitted from a high-power infrared LED light (850 nm, Conrad, Item No. 491248-62) placed behind the glass bulb. The eyes were recorded at 60 frames per second (fps). The eye movement recording and stimulus synchronization were achieved via the same custom written Python application for freely swimming fish recording. The eye tracking and pre-processing were done offline by a custom written Matlab script in the same way described in the section above.

### Stimulus design for behaviour experiments

Eight combinations of translational and rotational optic flow stimuli were used in the global motion decomposition experiments in freely swimming fish (Figure 6C), which includes the pure forward/backward translation stimuli, the pure CW/CCW rotational stimuli, and the 4 different mixtures of these pure translation/rotation stimuli. The design detail of these optic flow stimuli was described in a previous section (“Optic flow stimuli design”). The relative strength for the translation and rotation components in all mixed optic flow stimuli, *w_trans_* and *W_rot_*, were set to 1 and 0.67 respectively. All optic flow stimuli for OMR experiments were normalized by the maximum local movement speed to the range of 0-60 visual degrees/second. We also measured the baseline OMR and OKR activities as the swimming trajectory and eye movement when a static binary noise pattern was displayed.

As shown in Figure 6E, we designed 3 variants of translation stimuli to determine if and which visual features were involved in the conjugate eye movement observed in the optic flow decomposition experiment. The pure translation and “T+R” mixture optic flow stimuli in the OKR experiments in mounted fish are constructed in the exact same way as above, except the local motion speed in the optic flow stimuli was reduced to the range of 0-30 visual degrees/second to better trigger OKR behaviour. The slanted translation was constructed by rotating the pure translation 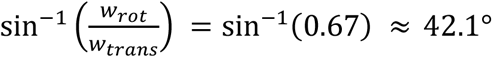, which corresponds to the azimuth of the contraction point of the “T+R” mixture. When the motion cue in one hemi visual field is closer than the other side, the speed difference is similar to the difference caused by the rotation interference. To mimic this, the asymmetric translation stimuli was constructed by multiplying each flow vector in the pure translation stimuli by the inverse of a distance factor (**D**) which was calculated as:

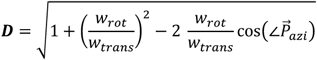

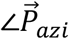 is the azimuth angle of the unit vector 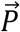 pointing from the observation point (the origin) to the location P in the visual field. The transformation mimicked the changes caused by shifting the observation points which are equidistant from all motion cues (***D*** = 1 a.u.) in the pure translation stimuli to the left by 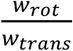 a.u.

### Retrograde labelling of nMLF

The retrograde labelling of the nucleus of the medial longitudinal fasciculus (nMLF) in nls-GCaMP6s zebrafish larvae was performed as described previously ^51^. Briefly, a 50% Texas Red dextran (10000MW, Thermo Fisher) in Evans solution was injected into the spinal cord of 4 dpf. larvae embedded in 1.7% agarose and anesthetized with tricaine. The injected fish was imaged with the two-photon microscope at 5 dpf.

## QUANTIFICATION AND STATISTICAL ANALYSIS

### Calcium data pre-processing

The midbrain and diencephalon of mounted fish were sampled in the calcium imaging recording in dorsoventral direction from 20 μm to 120 μm below the top of tectum with the step size 10 μm. The motion artifacts along the recording plane (XY plane) were corrected by a phase-correlation algorithm, and all visible neuron somata in the time-averaged image of the corrected recording video were automatically selected as ROIs by a marker-controlled watershed algorithm. Per-ROI calcium time series were extracted as the sum of all pixel values within the ROI for each frame.

A 3D z-stack that imaged along the dorsal-ventral axis from the top of tectum to deep ventral pretectum and dorsal thalamus (step size: 0.44 μm) was recorded for each fish. The ROIs in each recording were first registered to the corresponding z-stack. The neurons recorded in different fish were registered to the z stack of *Tg(elavl3:Hsa.H2B-GCaMP6s)mpn400* transgenic line provided by the Max Planck zebrafish brain atlas (mapzebrain) by registering the ROIs in each recording to the corresponding z-stack aligned with the atlas z-stack with a custom-written multimodal registration Matlab program.

### High throughput motion RF estimation

The high throughput motion RF estimation method used in our study was described in detail in a previous study (Zhang and Arrenberg, 2019). Briefly, the neural responses to the CMN stimulus were recorded with *in vivo* calcium imaging. As the abrupt, high amplitude increases of somatic calcium signals are usually correlated with significant increases of neural firing rate, these events were detected in the calcium traces and extracted for each ROI to obtain a binary event train (0 = no event, 1 = event detected). ROIs with event trains that only showed sporadic activity (#events/#stimulus frames < 8 %) were excluded for the following steps as we found only few of them could yield successful RF estimations with this method.

For each event train, an event triggered average (ETA) was computed as the raw RF estimation as follows:

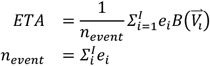

The e_i_ was the event state (0 or 1) for the i^th^ stimulus frame, and I is the total number of frames in the stimulus. To avoid the ambiguity in the vector summation, the corresponding motion vector field of the i^th^ frame 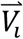 was transformed into a sparse, one-hot representation 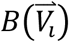 where the phase angles of motion vector are evenly divided into 16 direction bins as described in the previous study (see Zhang & Arrenberg, 2019 for details). For instance, for a vector 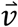 in 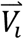 whose phase angle is in the range of the *k^th^* direction bin, its one hot transformation 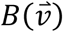 will be a 16-element array where the *k_th_* element is the vector norm 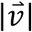 and the other elements are 0. Therefore, the size of these ETAs was 8 row×28 column×16 direction bins.

To identify the spatially connected ETA components significantly related with RF structures, a two-step non-parametrical cluster-based bootstrapping test (2-step NCB test) is applied. Briefly, the event train of each neuron was circularly permuted with random temporal offsets for 1000 times. By comparing the original ETA with the null ETAs computed from these permuted event trains, we determined an empirical probability for each unit in the original ETA. The original ETA units with an empirical probability value greater than 97.5% (their values are higher than 97.5% of the corresponding units in the null ETAs) were selected as potentially significant units (step I). To solve the multiple comparison problem in the detection of significant RF structures, we performed an additional cluster-based bootstrapping test (step II) where we grouped the selected ETA units in the original ETA connected in spatial and directional domain into significant clusters. We defined the cluster-level statistic as the sum of the empirical probability of all units in a significant cluster. By applying the same protocol to the null ETAs, we obtained a null cluster-level statistic distribution for each neuron. The clusters in the original ETA whose statistic were higher than 95% of the values in the corresponding null distribution are considered as significant RF components at the significance level of 5% (see the corresponding paper^28^ for detailed explanations). For the sake of clarity and convenience, the identified significant ETA components were converted from the one-hot representation form back to the vector field form for the downstream analyses and visualization. An RF mode is defined as a cluster of the spatially connected ETA components. The centres and preferred directions of RF modes were calculated as the averaged location and preferred direction of the components in each mode.

### Fitting optic flow fields to RFs

To determine the similarity of the estimated RF to translational or rotational optic flow fields, we identified the most similar translational and rotational optic flow field for each RF estimation by fitting the optic flow fields to RF estimations with the least squares estimation method. The residual sum of squared angular difference (*RSS_angle_*) between an RF estimation 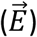 and the fitted optic flow field 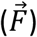 were calculated as:

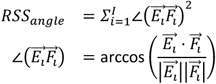

We solved the fitting problem by searching the translation/rotation axis that minimized the *RSS_angle_* with the Matlab function “*fminunc*”, a gradient descent optimizer. The similarity between the fitted translation/rotation flow fields and the motion RF estimations were calculated as the averaged cosine similarity between each vector pair of the RF-related ETA components and the optic flow fields as follows:

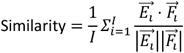

In addition, the symmetry of bimodal translation RFs in relation to the preferred translation directions in Figure 4E was calculated as:

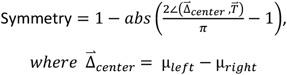

The μ*_left_* and μ*_right_* are the left and right mode center positions of the bimodal RFs. The bimodal RF is most symmetric when the vector linking its mode centers is perpendicular to its preferred translation direction 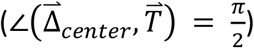.

### Prediction of optic flow preference

To determine how well the responses of corresponding neurons to optic flow can be explained by their RFs, we recorded the calcium responses of neurons in optic tectum and the pretectal areas in 3 fish to 36 global optic flow stimuli which include 6 pure translational and 6 pure rotational optic flow and the 24 combinations of these translational and rotational flow in random order (Figure 3E). Due to the memory limitation of our stimulus setup, we were unable to test all the combinations of translation and rotation optic flow fields, the 12 untested combinations were coloured in grey in Figure 3E. We chose these 12 combinations for exclusion, because their FoC/FoE positions are not shifted by the rotation components; we therefore expect that decomposition of rotation and translation is more easily accomplished for these combinations. The maximum speed for all optic flow stimuli were 30 degree/seconds. The translation and rotation component in the mixture optic flows were mixed in the ratio of 2:1 (*w_trans_* and *w_rot_*).

Each optic flow stimulus was displayed at 15 fps for 4 seconds followed with a 6-second pause before switching to next stimulus. The binary noise pattern of the last frame of the previous stimulus phase lasted over the pause period to become the first frame of the next stimulus phase, so the stimuli were switched seamlessly. After all optic flow stimuli had been displayed, the CMN stimulus was displayed for RF estimation in the same neurons.

The means and standard deviations of calcium fluorescence activities of each ROI in the 16-second period before and after all stimuli had been displayed were measured as μ_init_ / μ_end_ and σ_init_ / σ_end_. The ROIs whose |*μ_init_* - *μ_end_*| > 4*σ_init_* and/or *σ_init_/σ_end_* > 4 were excluded as either they might be bleached or drifted out of the imaged region. The calcium activities of the remaining ROIs were converted into z-score calcium traces as follows:

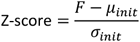

The z-score calcium traces (2 fps) were upsampled to match with the stimulus frame rate (15 fps). The response intensity (R) for each stimulus phase was calculated as:

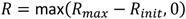

*R_max_* is the maximum normalized calcium activity (z-score) in a stimulus phase and *R_init_* corresponds to the normalized activity in the first frame of the stimulus phase (same as the last frame of the pause). Only the ROIs with estimated RFs ^28^ and strong responses to optic flow stimuli (*max*(*R*) > 3) were selected (1139 out of the 1484 ROIs with estimated RF in the experiments for Figure 3E-G). The RF-predicted responses (R_prediction_) to the presented optic flow stimuli 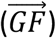 were calculated as follows:

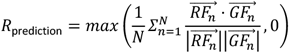

*N* is the number of significant components in the RF estimations, 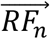 and 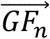 are the vectors of the n^th^ components in the RF and in the optic flow field respectively. Note in the following section, this function will be addressed as the “RF-based response function”.

The translation-rotation index (TR index) was computed as:

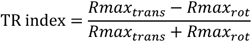

Rmax_trans_ and Rmax_ro_t are the maximum response intensity of all pure translation or rotation optic flow stimuli respectively.

### Translation direction and speed tuning

To determine the tuning properties of the translation-sensitive neurons found in the pretectal area, we measured their responses to the translational optic flows moving in 8 horizontal directions (azimuth: 0°,45°,90°,135°,180°,225°,270°,315°; elevation: 0°, Figure S4A) and 5 speed levels (7.5, 15, 30, 60, 120 degree/seconds). Each stimulus phase lasted for 15 seconds, including a 5-second moving phase and a 10-second pause phase. The stimuli were displayed in random order to avoid any historical effect.

The data pre-processing and the calculation of response intensity were the same as described in the previous section. We employed the discrimination index (DI) used in a previous study as a quantitative measure for the direction and speed tuning intensity separately ^52^:

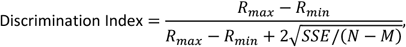

where *R_max_* and *R_min_* are the responses to the most and least effective stimulus, and the SSE is the sum squared error of the averaged responses, N and M are the number of trials and the degrees of freedom, respectively. Because the DI for one parameter of interest may be affected by the other irrelevant parameter, the *R_max_* and *R_min_* were calculated as the maximum and the minimum responses averaged for the irrelevant stimulus parameter. This way we could quantify DIs separately for speed and direction.

To quantitatively summarize the direction and speed tuning properties for all translation optic flow sensitive neurons, we fitted the joint distributions of von Mises and log Gaussian distributions to the neuron response data with the nonlinear optimization function “*fminsearch*” *in* MATLAB. The fitted distribution was:

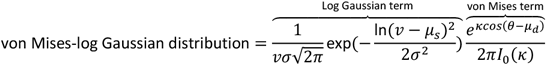

The log Gaussian and the von Mises term aimed to capture the speed and direction tuning properties, respectively. The preferred speed and direction of each neuron can therefore be quantified as the mode of the log Gaussian model *e^μ_s_-σ2^* and the von Mises model (*μ_d_*) respectively. The measure of concentration of the von Mises distribution K provides a quantitative measure of the direction tuning width. Note that the choice of log Gaussian function was mainly due to the non-negative property of speed and the similar shape to the speed tuning curves observed empirically. There was no biological basis for this model thus it was used only for description purpose.

### Population decoder for self-motion estimation

To determine if the information encoded by the translation-sensitive neurons was sufficient for determining the translation direction and speed, we constructed a population decoder derived from the maximum likelihood decoder with the calcium response data recorded. Since the accuracy of any decoding algorithm is limited under the Cramér Rao bound which is determined by the encoding information^53^, a low or no decoding error may prove the information encoded by these translation sensitive neurons is sufficient, even if the decoding algorithm implemented in zebrafish brain might be very different from our decoder.

As illustrated in Figure 5A (top), the ground truth stimulus parameter *T* is estimated as the least squares fit that minimizes the residual sum of squares between the calcium responses *R* of the recorded *n* translation-sensitive neurons, and the responses predicted by their response functions (*F*(*T*)):

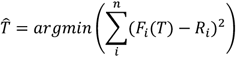

We computed the RF estimation of the neural responses to 1000 evenly sampled translation directions. The least squares fit, or the arguments of the minimum (argmin) is approximated by finding the nearest neighbour of the recorded responses in the sets of estimated responses with the Matlab function “*knnsearch*”.The histogram in Figure 5A contains 300 data points for “translation only”, which correspond to the six translation directions for which we had ground truth neuronal responses and 50 different bootstrapped decoders. Each decoder was constructed by drawing 50 random RFs out of the 138 translation RFs characterized in Figure 3E. The “mixture” histogram is based on 24×50 datapoints (24 global flow combinations from data in Figure 3E).

The responses functions were approximated with either the RF-based response (R_prediction_) function (Figure 5A bottom, see the STAR Methods section “Prediction of optic flow preference” for details) or the joint distribution model fitted to the direction-speed tuning data (Figure S4B-E, S5A-B). Since the decoding accuracy was limited by the encoding quality, the decoding error provides a quantitative measure for the encoding quality of translational optic flow. The error was calculated as the absolute difference between the real stimulus parameter and the estimation. In the decoder for translation and rotation mixtures (Figure 5A, bottom), the RF-based predictions to optic flow (from RF estimation using the CMN stimulus) were compared to the measured response to a particular optic flow (data from Figure 3E).

One problem in estimating the encoding quality of translation direction and speed using the joint distribution model in Figure S4B, is that the tuning model already contains the response information of the target stimuli to be tested with (which is different from the situation for the model at the bottom of Figure 5A). We employed a jack-knife test (“leave-ten-out”) to avoid this circular analysis: in the decoder for Figure S5A-B, the neural response data to translational optic flow with 40 different combinations of direction and speed parameters were evenly divided into 4 groups, so each group contained the responses to 10 different optic flow stimuli. For any target stimulus to be decoded, the decoder was only allowed to access the groups that did not contain the responses to the target stimulus for tuning model estimation.

To test the robustness of translation direction encoding algorithms implemented with the translation RFs, we constructed a population autoencoder which encodes the partially occluded or tampered translational optic flow field (used in Figure 5B and S5E) with *n* translation RFs into a one-dimensional array of neural responses as:

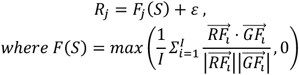

R_j_ and *F_j_*(*S*) are the simulated recorded response and the RF-based response function of the j^th^ simulated neurons to stimulus *S*, and *ε* is the noise term. The neural responses were decoded to estimate the translation direction with the same population decoder described above, which makes the decoding process basically a reverse operation of the encoding. Thus, the reconstructed error is representing mostly the encoding quality which is affected by the number of neurons involved (*n*), the noise in the encoding process (*ε*), and the input translational optic flow field 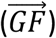.

To determine the noise effect on the encoding process, the reconstruction error of the autoencoder using all 329 translation RFs was tested with different encoding signal-to-noise ratio (SNR =+∞ (*ε* = 0), 2.6, 1.3 and 0.65) which was calculated as:

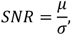

The μ is the averaged output from the response functions of all 329 simulated neurons, and σ is the standard deviation of the noise *ε*. For each SNR, the reconstruction error was tested for 100 random translation directions (Figure S5C).

We also evaluated the encoding quality for 100 random translation directions when the number of translation RFs involved *n* = 10, 20, 50, 100 and 329 (Fig. S5D, SNR = 1.3). Based on the results in Figure S5C-D, the autoencoder used in Figure 5B and S5E was constructed with 50 translation RFs and the SNR = 1.3. As a reference, on average, more than 100 translation sensitive neurons were found in the pretectal area per fish in the *in vivo* calcium imaging experiments, and as mentioned previously, the calcium signal SNR of the optic flow sensitive neurons analysed were all higher than 3.

The optic flow in naturalistic scenes is contiguous, so are the occlusions and object motions that partially block or tamper the optic flow fields. To mimic this contiguity, we created a 2D Gaussian blurred white noise matrix (σ= 12 visual degree) which had the same size as the optic flow fields for each optic flow field. Only the corresponding vectors in the optic flow field of the top N% elements (sorted by value) in the noise matrix were preserved in the final flow field (examples in Figure S5F). The rest of the optic flow field were occluded or replaced with random object motion (Figure 5B and S5E). For the latter, each spatially connected region in the rest of the optic flow field was assigned with a single random object motion. Similar to the calculation of translational optic flow fields, let the unit vector 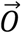 indicates the object motion in the 3D environment, the motion vector 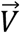 in a region of the visual field indicated by a unit vector 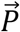 was calculated as:

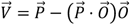

### Assessing vergence of optokinetic responses

A conjugate-vergence index (CV index) was computed for evaluating whether the OKR to each stimulus was more similar to a conjugate movement or a vergence movement:

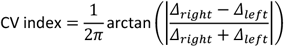

The CV index equals to 0 and 1 when the fish perform pure conjugate and vergence eye movement respectively. The CV indices of OKR to different optic flow stimuli were compared with one-sided, Bonferroni-corrected Wilcoxon rank-sum test to determine if any of them are significantly more similar to conjugate eye movement.

To determine if the conjugate eye movement to the rotation component in the T+R mixture optic flow is caused by local statistics, we randomly sampled the hemi optic flow fields for 1000 times and computed the averaged motion vectors for the left and right hemi field separately. The bootstrapped distribution of CV index in Figure 6F was computed in the same way from these pairs of bootstrapped motion vectors as described above.

## Supplementary videos

Video S1. The calcium activities of neurons in the pretectal area in response to pure translational optic flow, whose instantaneous direction is indicated by the spinning arrow, **related to Figure 4.**

Video S2. The calcium activities of the same neurons in Video S1 in response to mixed optic flow (translation + clockwise rotation), **related to Figure 4.** The instantaneous direction of the translation component is indicated by the spinning arrow.

Video S3. An example animation of a simulated fish (red) stabilizing its position in constant simulated “water current” by swimming against the estimated translational optic flow directions, **related to Figure 5.**

Video S4. An example clip of the OF stimulus (2D projected) used in Figure 4, **related to Figure 4.**

