## Supplementary figures for "A robust receptive field code for optic flow detection and decomposition during self-motion"

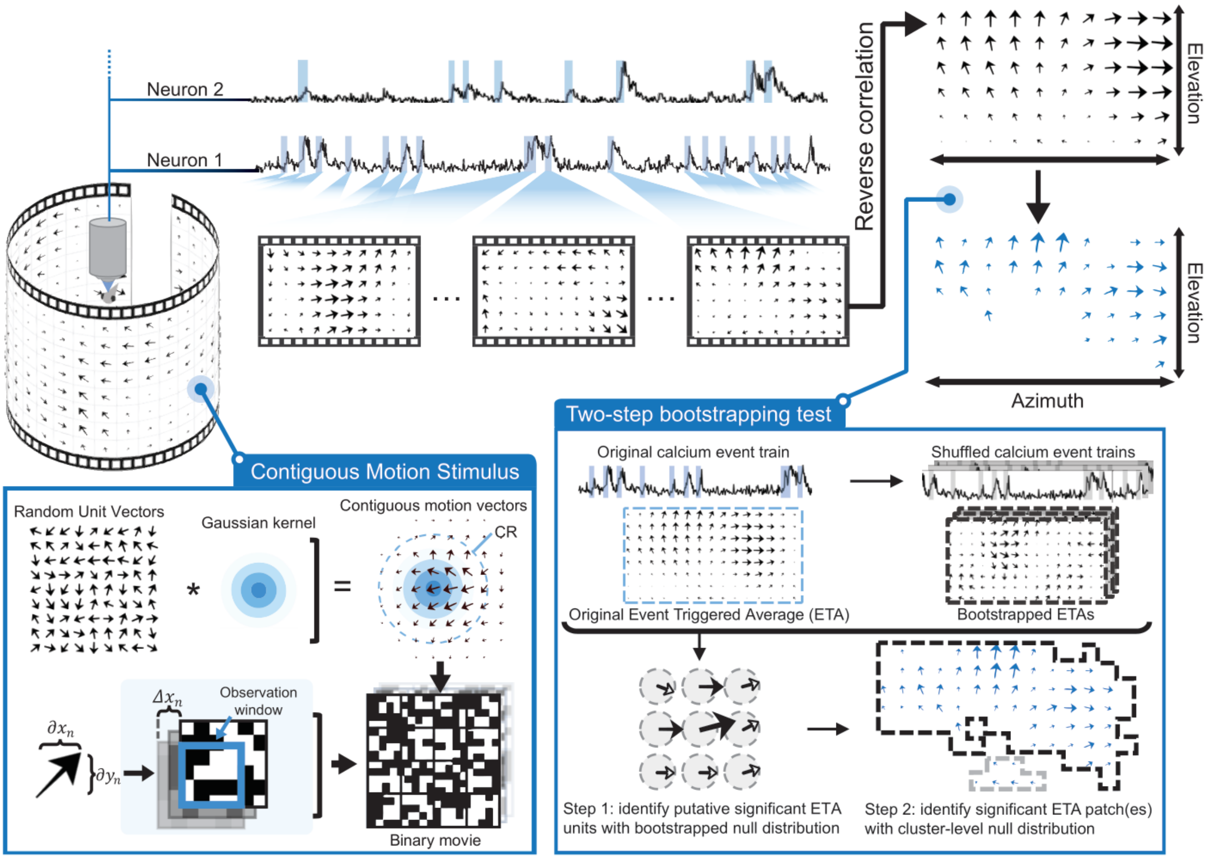


**Figure S1. A graphical abstract for the receptive field estimation method using contiguous motion stimulus and two-step bootstrapping test (adapted from Zhang and Arrenberg, 2019, see STAR Methods for details). Related to Figure 1B and STAR methods**.

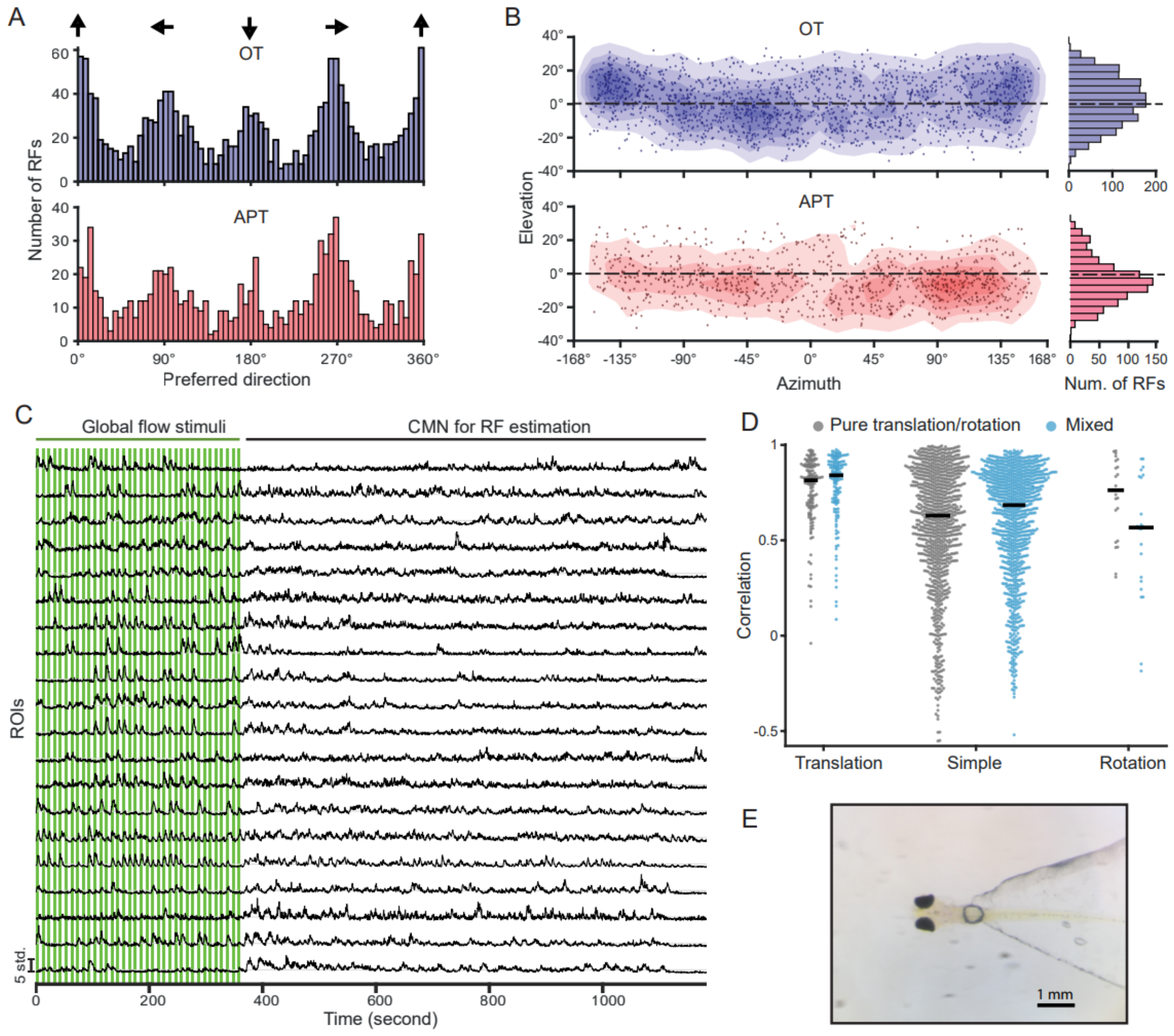


**Figure S2. Unimodal RF characterization and quantitative evaluation of complex RFs, related to Figure 1, 3 and STAR methods.** The distribution of averaged preferred directions (A) and RF centre locations (B) in the visual fields for the neurons with unimodal RFs in optic tectum (OT) and area pretectalis (APT). The black arrows in the top row of A indicate the preferred motion direction in 2D. Each dot in (B) indicates the centre of an RF on the 2D spherical coordinate map. The filled contours in the back of the scatter plot indicate the RF centre density estimated with kernel density estimation. The histograms on the right show the distribution of RF centre elevation. The RF centres of most unimodal neurons (66.8 %) in the APT but not the OT (45.8 %) were in the lower half of the visual field. (C) Examples of z-score calcium traces recorded in the experiment for Figure 3E-G. Each green shaded area indicates an optic flow stimulus phase (4 seconds duration + 6 seconds pause interval). (D) The bee-swarm plots of the correlation between the predicted and real neural response intensity to pure translational/rotational or mixed optic flows used in Figure 3E. The neurons were grouped by their RF types indicated in the x axis. (E) An image of an agarose-mounted 5 dpf zebrafish larva on the tip of a plastic triangle stage as used in this study.

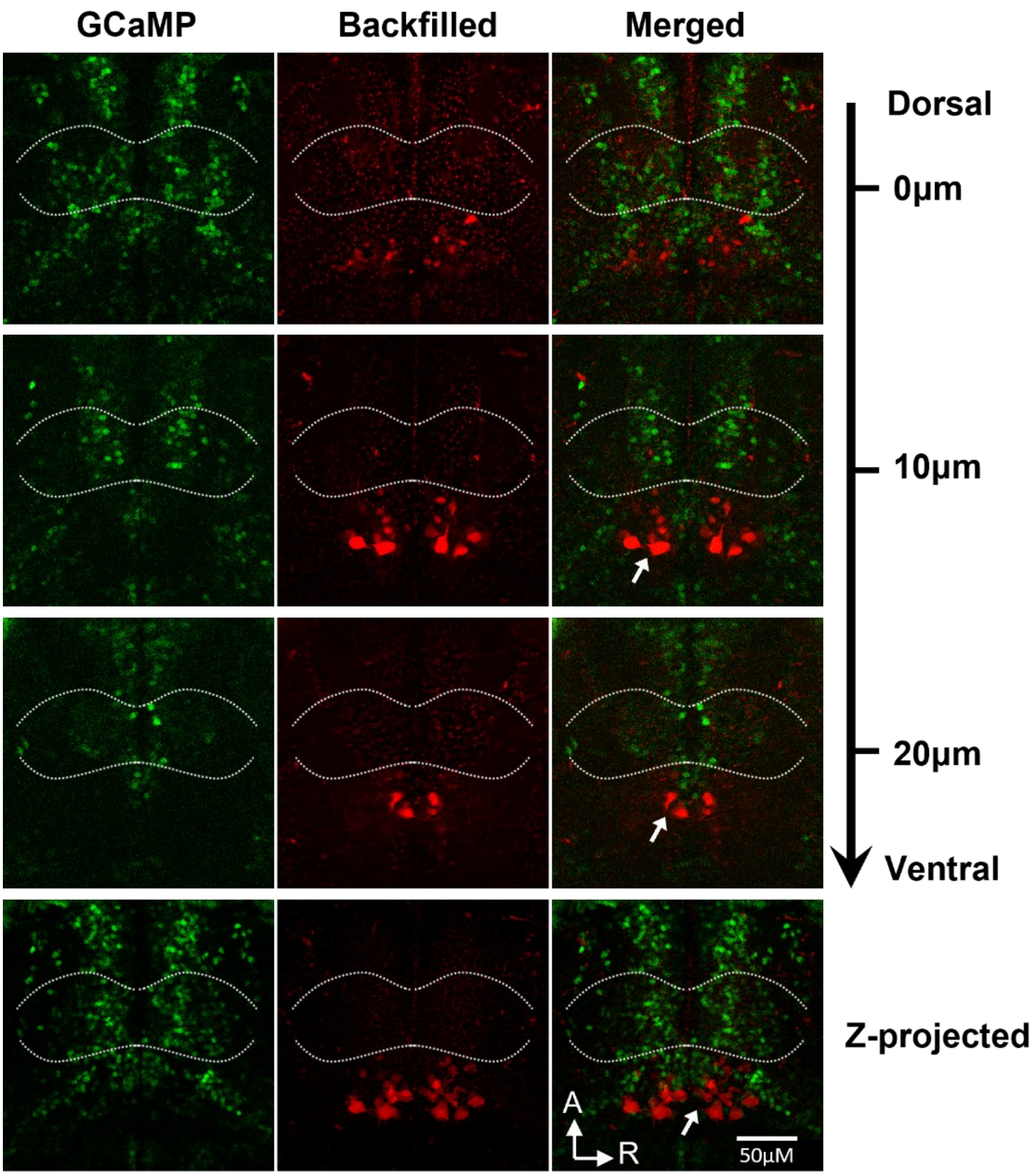


**Figure S3. Dorsal view of the pretectal area in 3 different depths (related to Figure 4B).** The green channel (GCaMP) displays all visible neural somata (two-photon calcium image) in the corresponding area, the white dashed curves outline the approximate location of the AMC pretectal clusters. The red channel shows the neural structures retrogradely labelled by spinal cord injections (see also STAR methods “Retrograde labelling of nMLF”). The white arrow in the merged channel indicates the nucleus of the medial longitudinal fasciculus (nMLF). The bottom rows contain the superimposed images of the pretectal area in different depth (Z-projected); A: anterior direction; R: right direction.

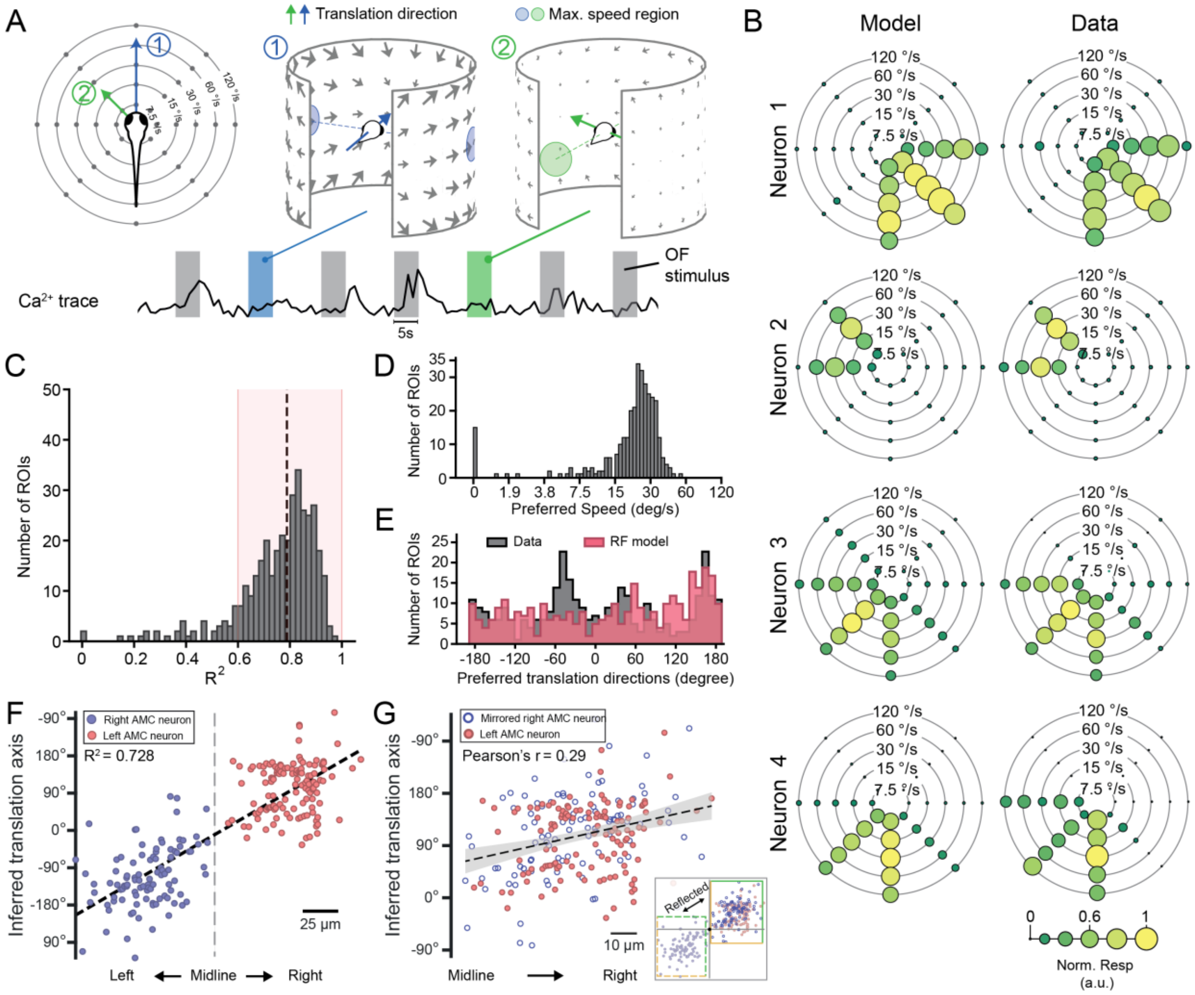


**Figure S4. Joint encoding of translational optic flow direction and speed in the pretectum, related to Figure 4.** (A) Left: Animals were presented with stimuli consisting of one of eight different TOF directions and five logarithmically spaced maximum velocities. Right: Two examples TOF stimuli are shown (① and ②). The region of maximum velocity of each stimulus is indicated as grey disk and located to the side of the translation direction (black arrow). To avoid the speed/distance ambiguity in TOF, which is not the focus of our study, all visual cues in our OF stimuli were equidistant to the animal. Therefore, the maximal angular speed in the translational optic flow is proportional to the fictive translation speed of the animal.(B) The recorded response (right column) and the fitted tuning model (left column, joint model of von-Mises and lognormal distributions)) for 4 example neurons. The response intensity is normalized by the averaged response amplitude for visualization purposes. (C) The $R^2$ distribution for the tuning model fitted to the neural responses to translational optic flow with different speed and/or direction. The black dashed line indicates the median $R^2$ and the red shaded area indicates the ROIs ($R^2$>0.6) used in (D-F) and Figure 4F.  (D) The preferred translation speeds of the translation-sensitive neurons inferred from the fitted tuning map. (E) The distribution of the preferred horizontal translation direction inferred from the fitted tuning model ("Data") and from the RFs of the neurons in the same anatomical locations ("RF model"). (F) Linear regression (black dashed line, $R^2$ = 0.728) between the preferred translation directions and the locations of AMC neurons on the left-right body axis. The dashed line indicates the midline of fish. (G) Intra-cluster linear correlation (Pearson's r = 0.29, p < 0.001) between preferred translation directions and the anatomical locations. The right AMC dots in (F) were reflected in the origin.

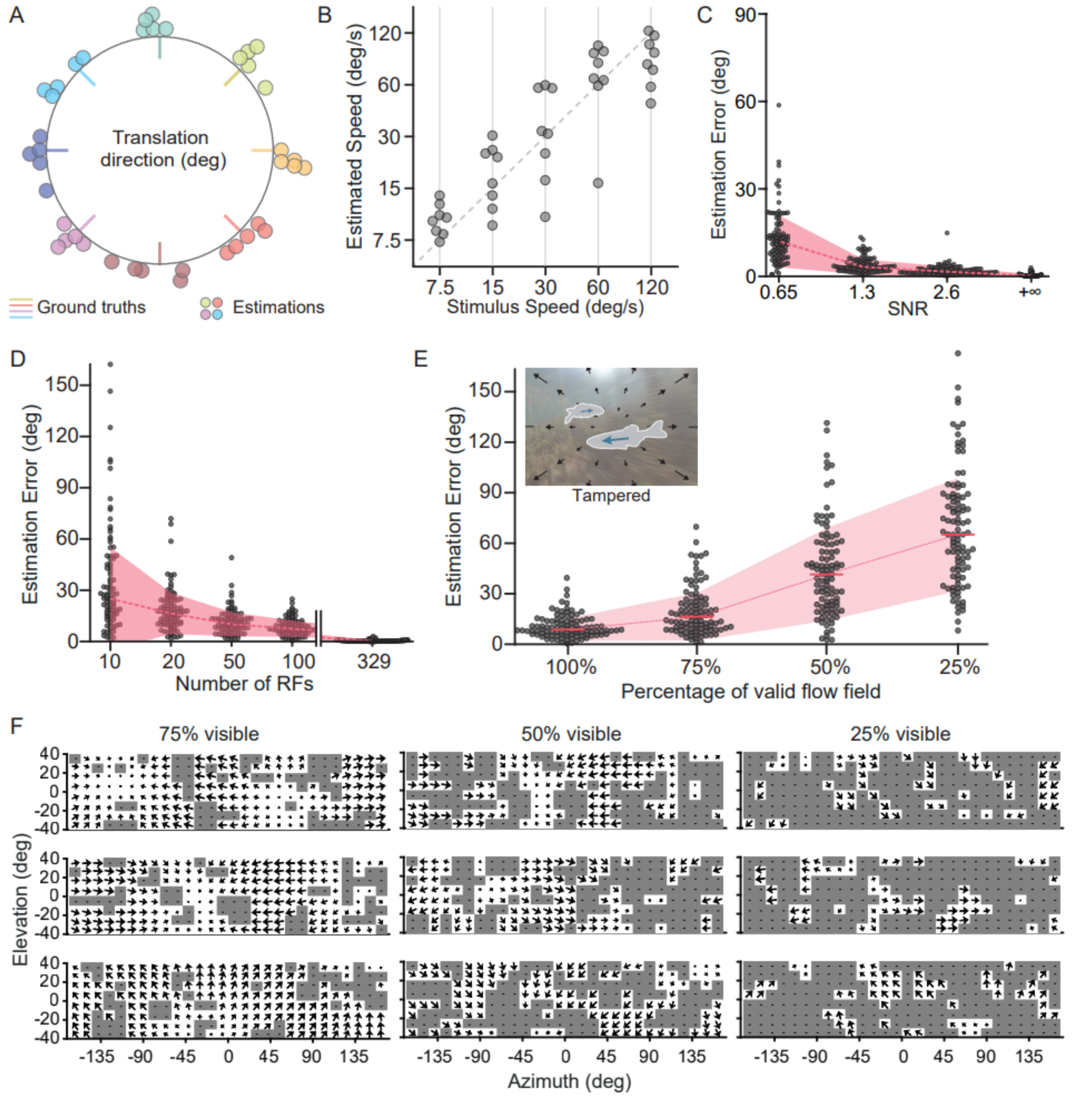


**Figure S5. Encoding accuracy of self-translation information in the population code of translation RF neurons, related to Figure 4, 5 and STAR methods.** (A) The translation direction estimated with the data presented in Figures 4F and S4A-E plotted in bee swarm style. Coloured bars indicate ground truth horizontal translation directions and circles the estimations. (B) The translation speed estimated with the data presented in Figures 4F and S4A-E in bee swarm style. The dashed line indicates the line of unity. (C-D) The bee-swarm plots of the estimation errors in relation to the signal-to-noise ratio (SNR) in the optic flow vector field (C) and the number of RFs contributing to the autoencoder (D). (E) Bee swarm plots of the estimation errors for the TOFs corrupted by moving objects. The dashed lines and shaded area in (C-E) indicate the median ± the standard deviation. (F) Examples of incomplete or tampered translational optic flow fields. Each grey block indicates a region in the visual field that contains no or wrong motion cue. In the tampered visual field, each set of spatially connected grey blocks displayed the 2D projected motion resulting from the 3D movement of a single object covering this region.

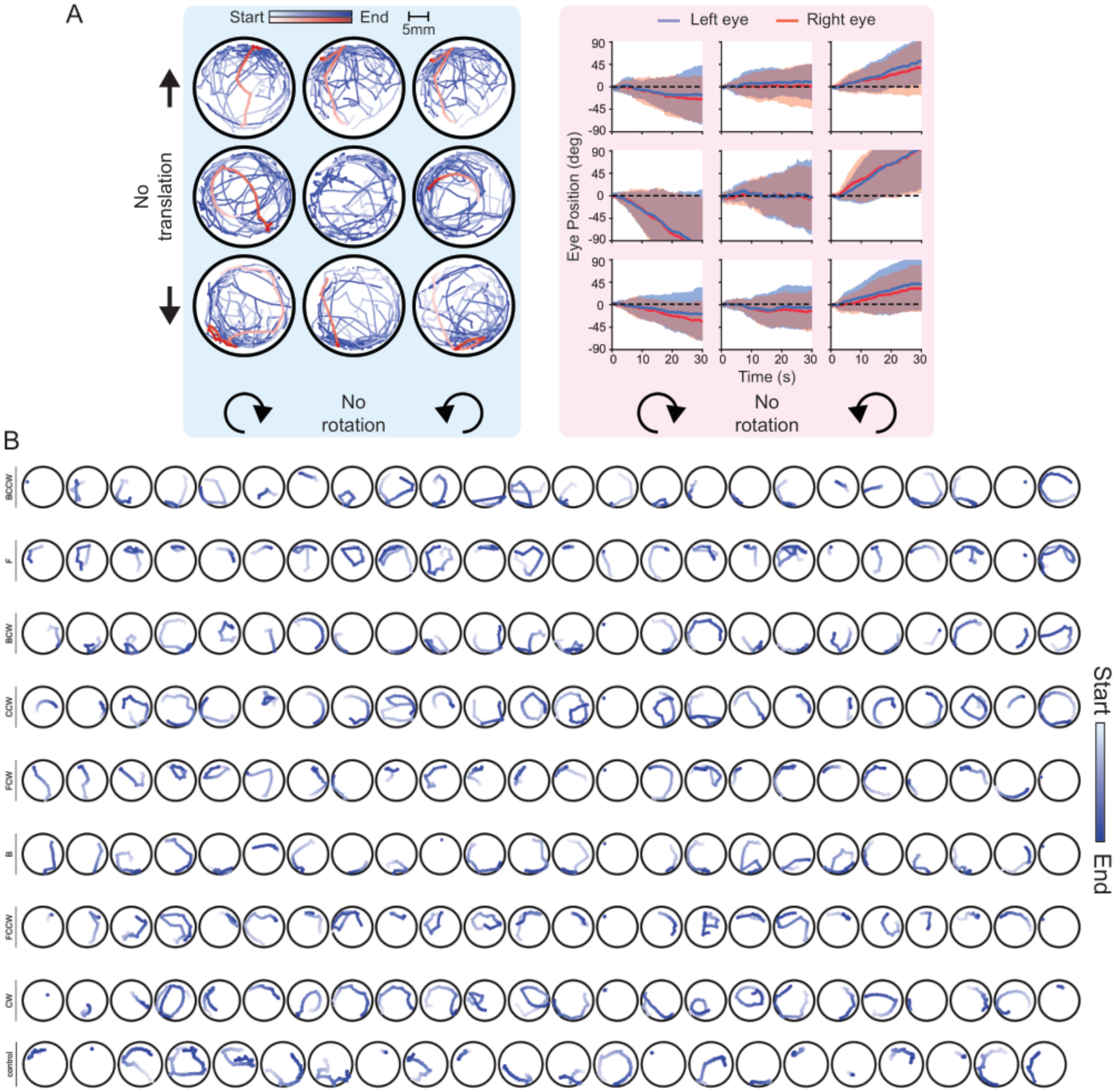


**Figure S6. OKR and OMR responses to all optic flow stimuli used (related to Figure 6).** (A) The superimposed swimming trajectories (left) during all 9 optic flow stimuli phases (indicated by the arrows at the sides) and the corresponding eye position traces (mean ± standard deviation). (B) The individual swimming trajectories for each optic flow stimulus (rows) and fish (column). From top to bottom: BCCW (backward translation + counter-clockwise rotation), F (forward translation), BCW (backward translation + clockwise rotation), CCW (counter-clockwise rotation), FCW (forward translation and clockwise rotation), B (backward translation), FCCW (forward translation + counter-clockwise rotation), CW (clockwise rotation) and control (stationary stimulus).
